# Functional metagenomics-guided discovery of potent Cas9 inhibitors in the human microbiome

**DOI:** 10.1101/569095

**Authors:** Kevin J. Forsberg, Ishan V. Bhatt, Danica T. Schmidtke, Barry L. Stoddard, Brett K. Kaiser, Harmit S. Malik

**Author notes:** Address correspondence to: Kevin J. Forsberg, 1100 Fairview Ave, A2-025, Seattle, WA 98109.

## Abstract

CRISPR-Cas systems protect bacteria and archaea from phages and other mobile genetic elements, which use small anti-CRISPR (Acr) proteins to overcome CRISPR-Cas immunity. Because they are difficult to identify, the natural diversity and impact of Acrs on microbial ecosystems is underappreciated. To overcome this discovery bottleneck, we developed a high-throughput functional selection that isolates *acr* genes based on their ability to inhibit CRISPR-Cas function. Using this selection, we discovered ten DNA fragments from human oral and fecal metagenomes that antagonize *Streptococcus pyogenes* Cas9 (SpyCas9). The most potent *acr* discovered, *acrIIA11*, was recovered from a *Lachnospiraceae* phage and is among the strongest known SpyCas9 inhibitors. *AcrIIA11* homologs are distributed across multiple bacterial phyla and many divergent homologs inhibit SpyCas9. We show that AcrIIA11 antagonizes SpyCas9 using a different mechanism than that of previously characterized inhibitors. Our study highlights the power of functional selections to uncover widespread Cas9 inhibitors within diverse microbiomes.

## Introduction

CRISPR-Cas adaptive immune systems are present in ~50% of bacterial and ~90% of archaeal genomes (Makarova et al., 2015), where they protect their hosts from infection by phages (Barrangou et al., 2007), plasmids (Marraffini and Sontheimer, 2008), and other mobile genetic elements (MGEs) (Zhang et al., 2013). CRISPR-Cas systems mediate this defense by incorporating short (~30 bp) spacer sequences from invading genomes into an immunity locus in the host genome. These spacer sequences are then expressed and processed into CRISPR RNAs (crRNAs) that, together with various Cas nucleases, mediate homology-dependent restriction of invading genomes. CRISPR-Cas systems are classified into six types (I, II, III, etc.) and 25 subtypes (I-A, I-B, II-C, etc.) on the basis of functional differences, phylogenetic relatedness, and locus organization (Makarova et al., 2018).

In response to restriction, phages and other MGEs have evolved dedicated CRISPR-Cas antagonists, called anti-CRISPRs (Acrs) (Bondy-Denomy et al., 2013), which can promote phage infection, enable horizontal gene transfer (HGT), and thus shape microbial ecosystems (Borges et al., 2017; Pawluk et al., 2017). Acrs that inhibit the type II Cas9 (Pawluk et al., 2016a; Rauch et al., 2017) and type V Cas12 systems (Marino et al., 2018; Watters et al., 2018) have also garnered significant interest in biotechnology, with demonstrated utility for reducing off-target Cas9 lesions (Shin et al., 2017), suppressing gene drives (Basgall et al., 2018), and precisely controlling synthetic gene circuits (Nakamura et al., 2019).

Work over the past decade has revealed much about the activity (Hille et al., 2018), distribution (Makarova et al., 2015), and evolution (Koonin and Makarova, 2017) of CRISPR-Cas systems. In contrast, comparatively little is known about Acr diversity and function. Known Acrs inhibit only seven of the 25 CRISPR-Cas subtypes (Bondy-Denomy et al., 2018; Makarova et al., 2018), though it is quite likely that unidentified Acrs antagonize the remaining 18 groups (Pawluk et al., 2017). In cases where antagonists of CRISPR-Cas systems have been identified, many additional undiscovered Acrs almost certainly exist that antagonize those systems (Watters et al., 2018). These undiscovered Acrs likely act via a diversity of mechanisms to influence gene flow and phage dynamics in microbial communities (van Belkum et al., 2015; Westra et al., 2016; Borges et al., 2017), and may unlock new modes of manipulating phage-and CRISPR-Cas-enabled technologies (Sheth et al., 2016; Knott and Doudna, 2018).

Previous efforts to discover *acr*s have relied on phage genetics (Bondy-Denomy et al., 2013; Pawluk et al., 2014; Hynes et al., 2017; He et al., 2018), linkage to conserved genes (Pawluk et al., 2016a; Pawluk et al., 2016b; Hynes et al., 2018; Lee et al., 2018; Marino et al., 2018) or the presence of a self-targeting CRISPR spacer, which would create an unstable autoimmune state if not for the presence of an Acr protein (Rauch et al., 2017; Watters et al., 2018). These clever strategies have revealed many candidate *acr*s. But they have also highlighted the difficulty of finding new *acr* genes based on homology, since *acrs* share little sequence conservation (Sontheimer and Davidson, 2017). As a result, most *acr*s almost certainly lie unrecognized among the many genes of unknown function in phages, plasmids, and other MGEs (Hatfull, 2015).

To overcome the challenges associated with anti-CRISPR discovery, we devised a functional metagenomic selection that identifies *acr* genes from any cloned DNA, based on their ability to protect a plasmid from CRISPR-Cas-mediated destruction. Because functional metagenomics selects for a function of interest from large clone libraries (Handelsman, 2004), it is well-suited to identify individual genes like *acr*s that have strong fitness impacts (Iqbal et al., 2014; Forsberg et al., 2015; Forsberg et al., 2016; Genee et al., 2016). This approach may be particularly useful for Acr discovery because Acrs are expressed from single genes and function readily in many genetic backgrounds (Pawluk et al., 2016a; Rauch et al., 2017).

Using this functional selection, we find that many unrelated metagenomic clones from human oral and gut microbiomes protect against *Streptococcus pyogenes* Cas9 (SpyCas9), the variant used most commonly for gene editing applications (Knott and Doudna, 2018). We identify a broadly distributed but previously undescribed Acr from the most potent Cas9-antagonizing clone in our libraries. This Acr, named AcrIIA11, binds both SpyCas9 and double-stranded DNA (dsDNA) and exhibits a novel mode of SpyCas9 antagonism, protecting both plasmids and phages from immune restriction. Thus, our functional approach reveals not only new classes of Acrs, but also new mechanisms of CRISPR-Cas antagonism.

## Results

### A functional metagenomic selection for type II-A anti-CRISPRs

We designed a functional selection to isolate rare *acr*s from complex metagenomic libraries. We based this selection on the ability of an *acr* gene product to protect a plasmid, which bears an antibiotic resistance gene, from being cleaved by SpyCas9 (figures 1A, S1). By screening metagenomes, our selection interrogates core bacterial genomes as well as DNA from the phages, plasmids, and other mobile genetic elements that infect these bacteria, which must contend with CRISPR-Cas immunity. Because most DNA inserts in large metagenomic libraries lack an *acr*, most clones in a library should be susceptible to SpyCas9-mediated destruction. However, those few DNA inserts that encode a functional Acr will resist SpyCas9 and can be recovered using the antibiotic resistance conferred by the plasmid they protect.

**Figure 1.**
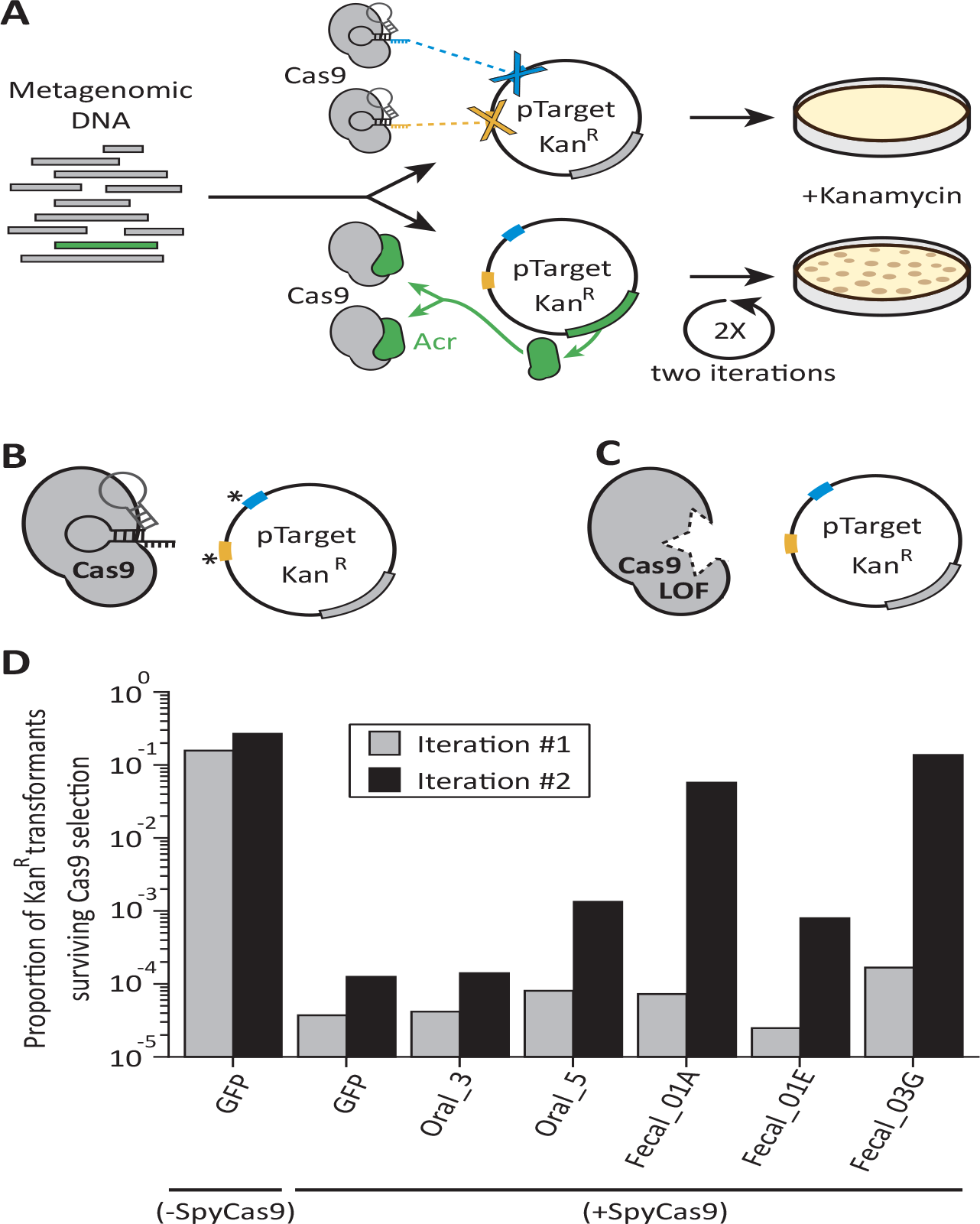
A functional metagenomic selection for type II-A anti-CRISPRs. **(A)** Plasmids in a metagenomic library without *acr*s are targeted by Cas9 and eliminated. Those few with *acr*s (green) withstand Cas9 and can be recovered via kanamycin selection. However, individual target site mutations, indicated by an asterisk **(B)** or Cas9 loss of function (LOF) mutations **(C)** allow plasmids to evade Cas9 independent of the metagenomic DNA insert that they carry. To reduce these major sources of false positives, we employ two target sites (blue, yellow) and two rounds of selection. **(D)** Relative to a GFP control, metagenomic libraries protect plasmids after two iterations through Cas9-inducing conditions. Each bar represents a single experiment.

In our *acr* selection scheme (figure S1), we targeted an inducible SpyCas9 nuclease to a kanamycin resistance (*Kan*^*R*^) gene on a plasmid used to construct the metagenomic libraries. Anticipating that resistance to Cas9 cleavage could arise readily via point mutations in target sites, we targeted Cas9 to two distinct loci within the *Kan*^*R*^ gene (figures 1B, S1) to reduce the frequency of these ‘escape’ plasmids. We transformed metagenomic libraries into an *Escherichia coli* strain that contained SpyCas9 under an arabinose-inducible promoter. After the cells were allowed to recover, we grew them overnight with arabinose to induce SpyCas9 expression. During this phase, cells were not exposed to kanamycin and thus were not under selection to maintain the metagenomic library bearing the *Kan*^*R*^ gene. We then plated cells on solid medium with kanamycin, killing cells in which SpyCas9 cleaved the *Kan*^*R*^ gene and allowing the recovery of metagenomic DNA from the few remaining Kan^R^ colonies.

Our analysis revealed that *Cas9* loss-of-function mutants (figure 1C) dominated the population in early, single-iteration experiments, occurring in approximately 10^−4^ to 10^−5^ transformants (table S1, figure S2). We therefore added a second iteration of SpyCas9 selection, reasoning that the Cas9 loss-of-function rate would remain constant across iterations, whereas *acr*-encoding clones would be enriched by a factor of ~10^4^ with each iteration. In theory, this enables us to select for an *acr* gene even if it is present just once in a library of ~10^7^ clones. To accomplish these two rounds of Cas9 selection, we purified total plasmid DNA from Kan^R^ colonies following one iteration, removed the original Cas9 expression plasmid via digestion, transformed the surviving plasmids into a fresh SpyCas9-expressing strain, and exposed them to SpyCas9 selection a second time (figure S1). As expected, adding a second iteration of Cas9 selection resulted in a significant enrichment for SpyCas9 antagonists above background (figure 1D).

### SpyCas9 antagonism in human oral and fecal metagenomes

We used our functional selection to search for *acr*s in five metagenomic libraries: two oral metagenomes from Yanomami Amerindians (Clemente et al., 2015) and three fecal DNA metagenomes from peri-urban residents of Lima, Peru (Pehrsson et al., 2016). We subjected each of these libraries, with an estimated 1.3×10^6^ - 3.4×10^6^ unique clones per library, to SpyCas9 selection (table S2). For each library, we observed a 10^4^ to 10^5^-fold reduction in the proportion of Kan^R^ colony forming units (CFU) following one iteration of SpyCas9 selection. This value matches the reduction in Kan^R^ CFU seen for a GFP control and the empirically determined frequency of Cas9 loss-of-function mutations (table S1, figures 1D, S2). Since Kan^R^ is a measure of plasmid retention, this result indicates that most clones in each library do not withstand SpyCas9. Following a second round of SpyCas9 selection for each library, metagenomic inserts were amplified from pooled Kan^R^ colonies by PCR, deep-sequenced, and assembled *de novo* using PARFuMS (Forsberg et al., 2012). Read coverage over each assembled contig was used to estimate its abundance following selection. After quality-filtering (*e.g.* removal of low-abundance contigs), we recovered a total of 51 contigs across all five libraries that putatively antagonize SpyCas9 (tables S3, S4, figure S3).

After two rounds of SpyCas9 selection, two libraries (Oral_5, Fecal_01A) seemed more likely to contain new *acr*s than the other three (Oral_3, Fecal_01E, Fecal_03G). The Oral_3 library poorly withstood SpyCas9 restriction, so was not studied further (figure 1D). While the Fecal_03G library completely resisted SpyCas9, subsequent analysis revealed that this was almost entirely due to a single clone that acquired mutations in both Cas9 target sites (table S3, figures 1B, S3). Accordingly, just one contig from this library passed quality filters. In contrast, the Fecal_01E library showed intermediate SpyCas9 resistance and was largely devoid of ‘escape’ mutations (figure 1D). However, just one of the 18 contigs from this library was found to be phage-associated (table S4). Because *acr*s are expected to originate in phages and MGEs (Borges et al., 2017; Pawluk et al., 2017; Sontheimer and Davidson, 2017), we predicted that contigs from Fecal_01E were unlikely to contain *bona fide acr*s and did not prioritize them in this study. These contigs may nonetheless encode anti-Cas9 activity, perhaps via Cas9 regulatory factors employed by host bacteria rather than MGEs (Hoyland-Kroghsbo et al., 2017; Faure et al., 2018), and therefore could represent a useful resource for probing host regulation of Cas9 activity.

Contigs from the Oral_5 and Fecal_01A libraries looked most promising for *acr* discovery. These libraries conferred SpyCas9 resistance at levels 10-fold to 1,000-fold above background (figure 1D) and had few escape mutations in Cas9 target sites, so likely withstood SpyCas9 due to functions encoded by their DNA inserts (table S3). Many contigs from these libraries are phage-associated (30%; table S4, figure 2A) and many encode genes of unknown function (figure S4); these are both hallmarks of known *acr*s (Borges et al., 2017; Pawluk et al., 2017; Sontheimer and Davidson, 2017). Additionally, we observed that Cas9 homologs are found in 15 of the 18 bacterial genera identified in these selections (83%, table S4), a significant enrichment over the 12% of bacterial genera that harbor Cas9 (567/4822) from a representative set of ~24,000 bacterial genomes (Mendler et al., 2018) (p=4×10^−21^, χ² test). This enrichment strongly supports our hypothesis that many of the recovered contigs encode true *acr*s rather than artifacts of functional selection, since phages and MGEs must encounter Cas9 to benefit from the protective effect of *acr*s.

**Figure 2.**
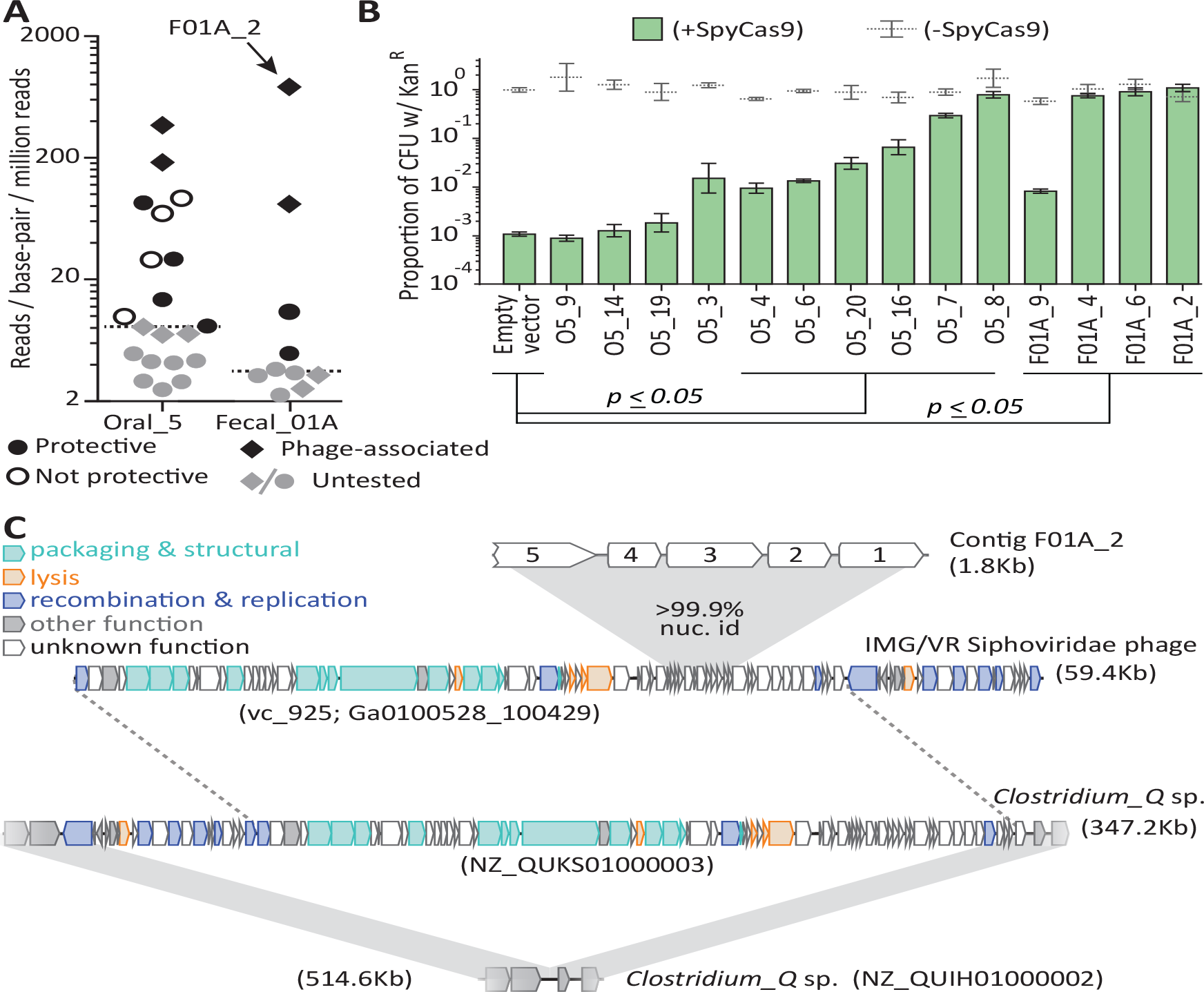
Metagenomic DNA inserts antagonize SpyCas9. **(A)** The read-coverage of all contigs from libraries Oral_5 and Fecal_01A is shown. Dashed lines depict median coverage for each library. Relative coverage can be used as a proxy for contig abundance within a library but cannot be used for meaningful comparisons across libraries. **(B)** Upon individual re-testing, six oral and four fecal inserts protect a plasmid from SpyCas9 relative to an empty vector (student’s t-test); +/- refers to SpyCas9 induction. Error bars depict standard error of the mean. **(C)** Contig F01A_2 is from a Siphoviridae phage which lysogenizes *Clostridium_Q* sp. Gray, shaded regions depict key regions of near perfect (>99%) nucleotide identity. Homology between the *Clostridium_Q* prophage and the IMG/VR phage is not depicted as these phages are nearly identical (except for one translocation or assembly-artefact involving the recombination and replication machinery). All accession numbers denote NCBI Genbank IDs except for the Siphoviridae phage, for which we use IMG/VR convention and indicate viral_cluster with the scaffold_id in parentheses. Contig lengths are depicted next to each sequence.

To confirm that the Oral_5 and Fecal_01A libraries encoded Acrs, we re-cloned the most abundant contigs from these libraries into fresh plasmid and strain backgrounds and tested them individually for SpyCas9 resistance. This step eliminated potential mutations to the plasmid backbone, the *cas9* gene, or the host genome that may have accounted for Cas9 resistance in the original screen. For 10 of 14 re-cloned contigs, the DNA insert still protected its parent plasmid from SpyCas9, confirming the power of our selection to identify novel *acr*s from metagenomic DNA (figures 2A, 2B). Intriguingly, none of the contigs recovered by functional selection encoded homologs of previously identified Acrs.

### Discovery of a widespread, potent Cas9 antagonist: AcrIIA11

Among the 10 contigs we confirmed as having Cas9 antagonist activity, several features made us focus on a single contig, F01A_2 (figure 2A). First, the F01A_2 contig was recovered from our iterative selection of the Fecal_01A library, which conferred near-complete protection against SpyCas9 (figure 1D). Second, the F01A_2 contig was by far the most abundant contig from this library (with coverage 216-fold above the median coverage in the library), suggesting that it outperformed other contigs during selection. In our re-testing, we confirmed that F01A_2 completely inhibited SpyCas9 activity (figure 2B). Finally, F01A_2 shares near-perfect nucleotide identity (>99.9%) with a *Siphoviridae* phage. This phage infects bacteria from the genus *Clostridium_Q* (figure 2C), within the family *Lachnospiraceae* (Parks et al., 2018). As is typical for *Acr*-encoding loci, the five open reading frames (ORFs) on F01A_2 are small, map to accessory regions of phage genomes, and appear to routinely undergo HGT (figure S5).

To identify the ORF(s) in F01A_2 responsible for SpyCas9 antagonism, we introduced an early stop codon into each of the five predicted ORFs in the contig and re-tested this set of null mutants for anti-Cas9 activity. *Orf_3* completely accounted for SpyCas9 inhibition: a null mutation in *orf_3* reduced the frequency of Kan^R^ 10^5^-fold, to the level of an empty-vector control (figure 3A). Furthermore, *orf_3* was sufficient for SpyCas9 antagonism, protecting a target plasmid (figure 3B) from SpyCas9 approximately as well as *acrIIA4*, the most potent inhibitor of SpyCas9 yet described (Rauch et al., 2017).

**Figure 3.**
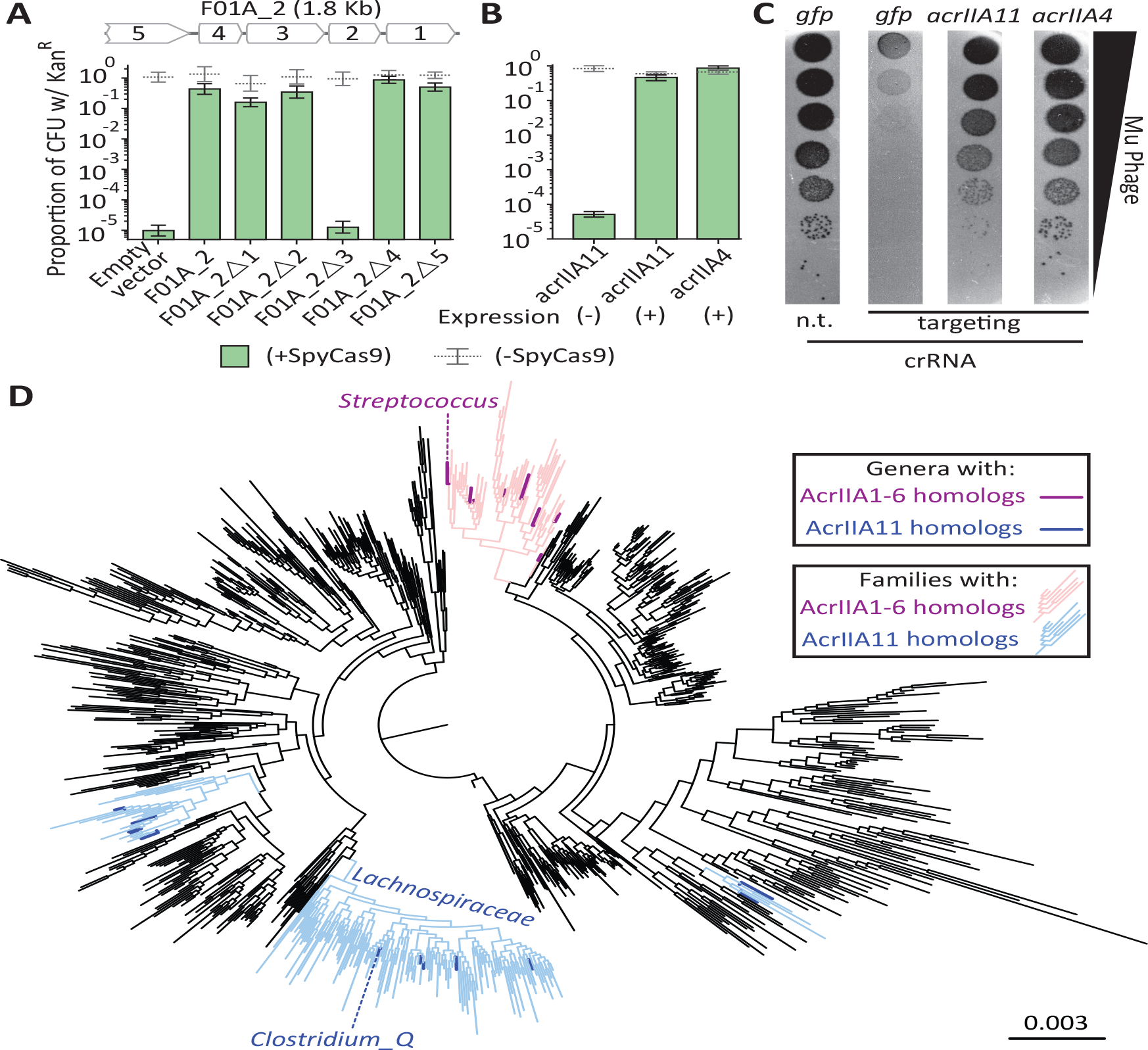
AcrIIA11 is a widely disseminated anti-CRISPR that protects plasmid and phage from SpyCas9. **(A)** The F01A_2 contig is depicted above the bar chart. Delta symbols (Δ) indicate early stop codons in each gene of the contig. Only the third gene on contig F01A_2 is necessary for SpyCas9 antagonism. **(B)** Induction of the third gene, named *acrIIA11*, is sufficient for SpyCas9 antagonism, protecting a plasmid as well as *acrIIA4*. Error bars depict standard error of the mean. **(C)** Mu phage fitness, measured by plaquing on *E. coli* expressing Mu-targeting SpyCas9, is measured in the presence of *gfp*, *acrIIA11*, or *acrIIA4* via serial ten-fold dilutions (also see replicated in Figure S6). Based on a non-targeting (n.t.) crRNA control, we conclude that SpyCas9 confers 10^5^-fold protection against phage Mu in these conditions. Both *acrIIA11* and *acrIIA4* significantly enhance Mu fitness by inhibiting SpyCas9. **(D)** A phylogenetic tree of Firmicutes and related phyla that shows the widespread dissemination of *acrIIA11* relative to other type II-A *acr*s. Each node represents a genus; families colored light blue contain *acrIIA11* homologs while families colored pink contain homologs of *acrIIA1 - acrIIA6*. Genera are colored dark blue or magenta if *acr* homologs could confidently be assigned to this phylogenetic resolution. The phylogenetic placement of *Clostridium_Q* and *Streptococcus* is shown. See figure S7 for a completely annotated version of this phylogeny.

We also investigated the ability of *orf_3* to restore phage infection in the face of SpyCas9-mediated immunity. We tested SpyCas9’s ability to prevent Mu phage infection in the absence of any Acr or in the presence of either *acrIIA4* or *orf_3*. Like with the plasmid protection assay, we found that *orf_3* provides phage with nearly complete protection from SpyCas9, similar to *acrIIA4* (figures 3C, S6). Given its origin from a phage contig, and its ability to potently inhibit SpyCas9 in both plasmid and phage protection assays, we renamed *orf_3* from contig F01A_2 as *acrIIA11*, to distinguish it from *acrIIA4* and other previously identified *acr*s (Borges et al., 2017; Pawluk et al., 2017; Lee et al., 2018; Uribe et al., 2019), to which it bears no discernible homology.

Since *acrIIA11* is a newly discovered *acr*, we wished to determine its distribution in nature. We therefore searched for homologs in NCBI and in IMG/VR, a curated database of cultured and uncultured DNA viruses (Paez-Espino et al., 2017). We identified many proteins homologous to AcrIIA11 in both phage and bacterial genomes and focused on a high-confidence set of homologs, those with ≥35% amino acid identity over ≥75% of AcrIIA11’s 182 amino acid sequence. AcrIIA11 homologs have a wider phylogenetic distribution than most previously identified type II-A anti-CRISPRs (figures 3D, S7) and span multiple bacterial phyla. We made a phylogenetic tree of AcrIIA11 homologs and found that they clustered into three monophyletic clades (figure 4A), which correspond to the three bacterial taxonomic groups in which they are found (figures 3D, S7). This concordance between gene and species clusters indicates that, while HGT of *acrIIA11* may routinely occur across short phylogenetic distances (figure S5), intra-class and intra-phylum gene flow is rare, in contrast to what has been observed for some other *acr*s (Uribe et al., 2019).

**Figure 4.**
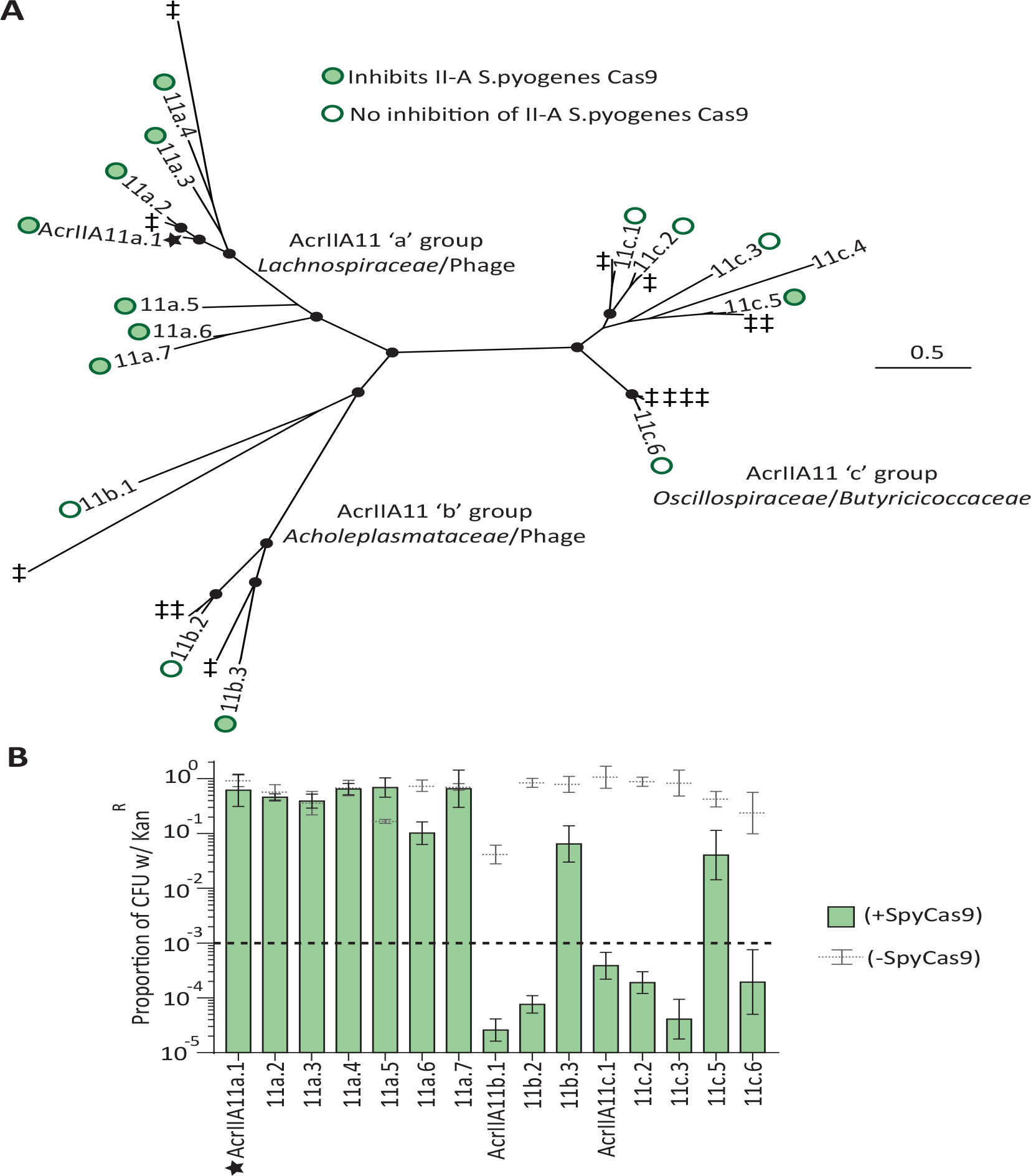
Diverse AcrIIA11 homologs inhibit SpyCas9. **(A)** An unrooted phylogenetic tree of predicted AcrIIA11 homologs clusters into three major clades that correspond to the bacterial families depicted in figures 3D and S7. Circles at nodes indicate bootstrap support >0.75. The star indicates the original AcrIIA11 sequence isolated via functional selection. Open and filled circles indicate whether a given homolog inhibits SpyCas9 in a plasmid protection assay. Untested homologs are indicated with a diesis (‡); toxicity with AcrIIA11c.4 prevented it from yielding meaningful data **(B)** AcrIIA11 homologs antagonize SpyCas9. The dashed lined indicates the threshold used to assign inhibitory labels in (A). Error bars depict standard error of the mean.

AcrIIA11 appears restricted to specific bacterial taxa that are distantly related to *Streptococcus*. Nevertheless, AcrIIA11 potently inhibits SpyCas9, even though SpyCas9 is highly diverged from the type II-A Cas9 proteins found in AcrIIA11-containing taxa. For instance, the only Cas9 protein in *Clostridium_Q* (NCBI accession CDD37961), AcrIIA11’s genus-of-origin, shares just 32% amino acid identity with SpyCas9. Intrigued by this observation, we tested divergent AcrIIA11 homologs from all three phylogenetic groups against SpyCas9 and found that homologs from each group could inhibit its activity (figure 4B). Type II-A CRISPR-Cas systems are enriched in the *Lachnospiraceae* relative to other bacterial families (figure S7, p=5×10^−27^, χ² test), which may explain why all tested AcrIIA11 homologs in the ‘a’ group (figure 4A) inhibited SpyCas9 (figure 4B). Based on our findings, we hypothesize that AcrIIA11 homologs may intrinsically possess the capacity to antagonize a diverse set of Cas9 orthologs. Because homologs of AcrIIA11 are found in many genera prevalent within human gut microbiomes (figure S7), its broad predicted range suggests that AcrIIA11 homologs will pose a meaningful barrier to Cas9 activity in this habitat – both in the context of natural phage infections as well as Cas9-based interventions to manipulate microbiome composition (Sheth et al., 2016; Pursey et al., 2018).

### A novel mode of SpyCas9 antagonism by AcrIIA11

Considering their relatively recent discovery, it is not surprising that very few Acrs have been mechanistically characterized. Nonetheless, a common theme has emerged from biochemical studies of AcrIIA2 and AcrIIA4, the only type II-A Acrs for which a mechanism of Cas9 inhibition has been elucidated. These studies have shown that both AcrIIA2 and AcrIIA4 are dsDNA mimics; they inhibit SpyCas9 by binding to its gRNA-loaded form and preventing association with a dsDNA target (Dong et al., 2017; Shin et al., 2017; Yang and Patel, 2017; Jiang et al., 2019; Liu et al., 2019). Given that AcrIIA11 is an unrelated inhibitor, we sought to address whether it has the same mode of SpyCas9 antagonism as these previously-studied type II-A Acr proteins.

We therefore purified recombinant AcrIIA11 from *E. coli* to analyze its biochemistry, interactions with SpyCas9, and possible modes of inhibition. We first observed that untagged AcrIIA11 migrates at approximately twice its predicted molecular weight by size exclusion chromatography (figure 5A), suggesting that it may function naturally as a dimer. Validating our genetic findings, purified AcrIIA11 inhibited the dsDNA cleavage activity of SpyCas9 in a concentration-dependent manner (figures 5B and 5C). These data demonstrate that AcrIIA11 is sufficient for SpyCas9 inhibition and does not require additional host factors. This autonomous activity is consistent with *acrIIA11*’s phylogenomic signature, as it shows no linkage to any other gene across phage genomes (figure S5).

**Figure 5.**
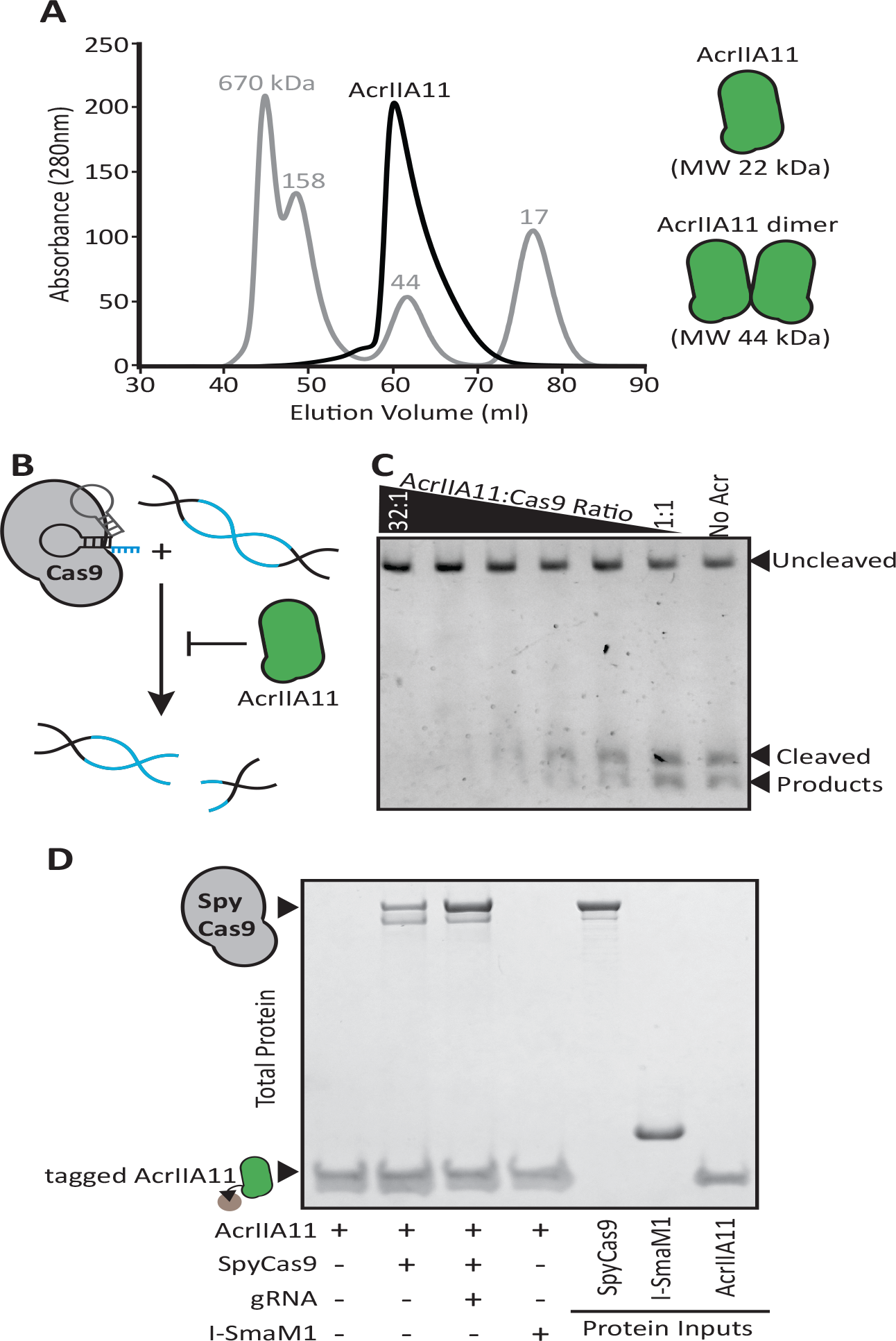
AcrIIA11 binds SpyCas9 to inhibit dsDNA cleavage. **(A)** AcrIIA11 elutes as a dimer on size exclusion chromatography. The gray trace depicts protein standards of the indicated molecular weight. The black trace shows AcrIIA11 elution. The predicted molecular weights of its monomeric and dimeric forms are depicted to the right. **(B)** Schematic of dsDNA cleavage assay in (C). **(C)** AcrIIA11 inhibits the ability of SpyCas9 (0.4µM) to cleave a linear 2.6 Kb dsDNA substrate in a concentration-dependent manner. **(D)** AcrIIA11 binds SpyCas9, with a moderate preference for the gRNA-loaded form. A Coomassie stain of total protein following pulldown of a 2x-strep-tagged AcrIIA11 incubated with either purified SpyCas9 (+/- gRNA) or the meganuclease I-SmaMI (as a negative control). SpyCas9 often runs as a doublet (see the protein marker), likely due to partial protein degradation.

We considered several possibilities for AcrIIA11’s mode of antagonism. We first tested whether AcrIIA11 prevents SpyCas9 from binding gRNA. We found that AcrIIA11 has no clear effect on gRNA binding, though it does bind gRNA at a large (160-fold) molar excess and may interact with the SpyCas9/gRNA complex (figure S8). We therefore tested whether AcrIIA11 directly binds SpyCas9. We found that AcrIIA11 was unable to bind the I-SmaMI meganuclease (used as a negative control) but bound both the apo (without gRNA) and gRNA-loaded forms of SpyCas9. The addition of gRNA enhanced AcrIA11 binding to SpyCas9 (figures 5D, S9) but not to the same extent as with AcrIIA4 binding SpyCas9 (figure S9). Strong binding to the gRNA-loaded form of SpyCas9 has been previously demonstrated for both AcrIIA2 and AcrIIA4 (Dong et al., 2017; Shin et al., 2017; Yang and Patel, 2017; Jiang et al., 2019; Liu et al., 2019).

Given that AcrIIA11 has similar Cas9-binding properties as AcrIIA2 and AcrIIA4, we next tested whether AcrIIA11 also acts as a DNA mimic to antagonize SpyCas9. Both AcrIIA2 and AcrIIA4 have the key property of preventing gRNA-complexed SpyCas9 from binding target dsDNA (Dong et al., 2017; Shin et al., 2017; Yang and Patel, 2017; Jiang et al., 2019; Liu et al., 2019). To test whether AcrIIA11 functions similarly, we performed an electrophoretic mobility shift assay (EMSA). Consistent with previous work (Dong et al., 2017; Shin et al., 2017; Yang and Patel, 2017), we found that AcrIIA4 prevents the gel-shift caused by SpyCas9/gRNA binding to its target dsDNA. In contrast, we observe that AcrIIA11 does not prevent this gel-shift, even at molar ratios that abolish SpyCas9 cleavage activity (compare lanes 2-5, figure 6A).

**Figure 6.**
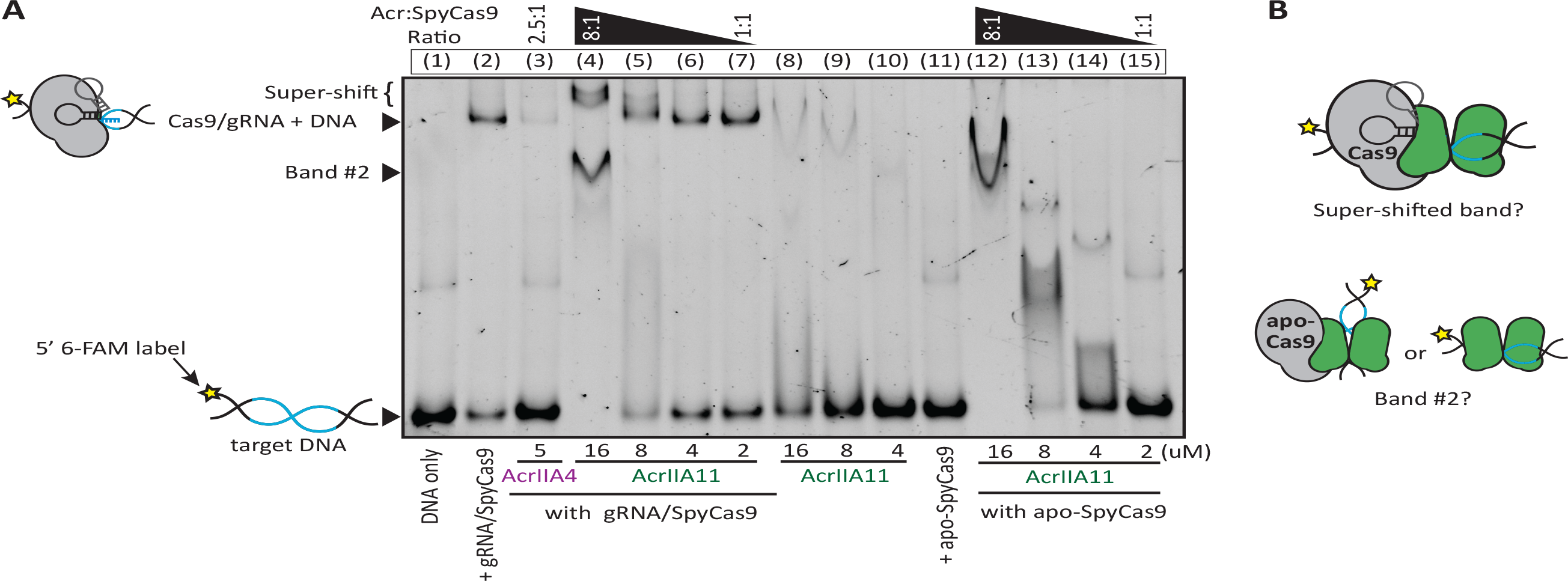
AcrIIA11 inhibits SpyCas9 via a novel mechanism. **(A)** Using an EMSA, we examine the gel shift experienced by target DNA (fluorescently labeled with a 6-FAM marker) incubated with either purified SpyCas9, gRNA, and/or purified Acrs. Divalent cations were omitted from reactions such that SpyCas9 could bind, but not cleave, target DNA, as previously shown (Lee et al., 2018). Lane numbers and Acr:SpyCas9 ratios are depicted above the native gel. Lane contents and Acr concentrations are depicted below the gel; 2uM SpyCas9 was used. Key bands are annotated to the left of the gel. **(B)** Schematics of possible explanations for the super-shifted band and band #2 in (A) are depicted.

Instead of preventing a SpyCas9-induced gel-shift, we see that AcrIIA11 creates the opposite effect, giving rise to a super-shifted SpyCas9/gRNA/dsDNA ternary complex (lane 4, figure 6A). This super-shift may result from AcrIIA11 binding the ternary complex (as in figure 6B). Alternatively, it could result from AcrIIA11 binding dsDNA overhangs unprotected by SpyCas9’s footprint. Indeed, we find that AcrIIA11 weakly binds DNA (lanes 8 and 9, figure 6A) but that DNA-binding is greatly enhanced in the presence of SpyCas9 (compare ‘band #2’ in lanes 4 and 12 with 8 and 11, figure 6A). Thus, AcrIIA11 and SpyCas9 could act together to bind DNA or one protein could stimulate the other’s DNA-binding ability (figure 6B). Although it remains unclear whether AcrIIA11’s mode of antagonism is related to the ‘super-shift’ we observe, our results nevertheless confirm that AcrIIA11 uses a different mechanism for SpyCas9 inhibition than AcrIIA2’s or AcrIIA4’s DNA mimicry.

## Discussion

The identification and characterization of CRISPR-Cas systems has outpaced the discovery of anti-CRISPRs, even though Acr diversity almost certainly exceeds CRISPR-Cas diversity in nature (Pawluk et al., 2017; Watters et al., 2018). It seems likely that these undiscovered Acrs are important influences on both phage infection outcomes and horizontal gene transfer in natural microbial communities (Borges et al., 2017). Determining the impact of Acrs on these processes is a challenge, however, because *acr*s lack shared sequence features so are difficult to detect. To overcome this obstacle and enable function-based *acr* discovery, we developed a high-throughput selection that identifies *acr*s based solely on their activity. Applying our functional selection to human fecal and oral metagenomes, we recovered 51 DNA fragments that putatively antagonize SpyCas9, with 10 confirmed for activity and many additional host-or phage-encoded inhibitors expected among the remaining 41 sequences.

In parallel work to ours, Uribe and colleagues recently described a functional metagenomic scheme to identify *acr*s (Uribe et al., 2019). They used a single round of Cas9 selection and one targeting crRNA to identify putative SpyCas9-antagonizing clones, though the authors suspected many of these to be false positives. To find *acr*s, they selected 39 ideal candidates from their initial dataset for re-testing, confirming 11 DNA fragments for anti-Cas9 activity, from which they identified four new *acr*s (*acrIIA7*-*10*). Although the basic premise of both of our approaches is similar, our strategy uses two iterations of Cas9 selection and two targeting crRNAs to identify *acr*s, which suppresses both Cas9 loss-of-function mutations and escape mutations in the target plasmid. As a result, we confirmed that 10 of the 14 most abundant clones following selection could antagonize SpyCas9 (figure 2B). More generally, we show that iterative selection can enable high-confidence discoveries from high-diversity inputs, which will be critical in the search for even rarer classes of *acr*s and other phage counter-defense strategies.

One of the major surprises from our work and that of Uribe *et al* is that most *acr*s in nature have not been discovered. Neither study encountered a previously described *acr* and no *acr*s were shared across datasets, though more than 10^7^ unique clones were screened for antagonists of the same Cas9 allele (SpyCas9) in the same selection host (*E. coli*). This lack of overlap emphasizes that the few *acr*s which have been catalogued are a very minor subset of those that are found in nature. This tremendous diversity exists because selection favors a highly-varied repertoire of *acr*s among even related MGEs (Bondy-Denomy et al., 2013; Borges et al., 2017) and Acrs have evolved independently many times from a variety of progenitor proteins (Rollins et al., 2018; Stone et al., 2018; Uribe et al., 2019). With extreme diversity favored, and the relative ease by which new Acr function is born, it is almost certain that much more mechanistic variety among Acrs remains to be discovered.

In this study, we have focused on the most potent SpyCas9 antagonist from our set, extensively characterizing the phylogenetic distribution, functional range, and mechanism of inhibition for its causal Acr, AcrIIA11. SpyCas9 is unlikely to be the natural target of AcrIIA11, as it is quite divergent from the Cas9 homologs found in AcrIIA11-encoding bacteria (Chylinski et al., 2014). Yet, diverse AcrIIA11 homologs, which are distributed across multiple phyla, retain inhibitory activity against SpyCas9. This suggests that AcrIIA11 is intrinsically predisposed to act against a wide variety of Cas9 sequences. Such inhibitory breadth may be particularly useful to MGEs that infect bacteria with diverse CRISPR-Cas systems (Makarova et al., 2015; Crawley et al., 2018) and could explain AcrIIA11’s broad phylogenetic distribution. This combination of prevalence and inhibitory breadth could also impact many Cas9-based interventions (Sheth et al., 2016; Pursey et al., 2018), as a pre-existing set of Acrs could quickly overcome diverse Cas9 homologs given the appropriate selective pressures. For instance, Cas9 inhibitors offer a means to resist Cas9 gene drives, so will have high selection coefficients if encountered during a gene drive (Bull and Malik, 2017). Gene drives, therefore, may inadvertently select for Acrs, especially those already abundant in microbiomes and with a high proclivity for HGT (Maxwell, 2016).

Broad inhibitory activity implies that AcrIIA11 might interact with conserved residues on Cas9. AcrIIC1, a broad-spectrum inhibitor of type II-C Cas9 enzymes, binds to conserved Cas9 residues to prevent nuclease activity but not target DNA binding (Harrington et al., 2017). No type II-A Cas9 inhibitor has been shown to act similarly, though AcrIIA11’s behavior is consistent with such a mechanism; it binds SpyCas9, inhibits DNA cleavage, but does not prevent target recognition. Alternatively, AcrIIA11’s broad predicted range may result from an indirect mode of inhibition that Cas9 cannot trivially escape. Such a mechanism could be related to AcrIIA11’s gRNA-binding ability, its DNA-binding capacity, or due to an interaction with SpyCas9 that alters these properties in either protein. While AcrIIA11’s exact mechanism still awaits elucidation, its mode of antagonism is clearly distinct from the other two mechanistically-characterized type II-A Acrs, AcrIIA2 and AcrIIA4, which act as dsDNA mimics to prevent target recognition (Dong et al., 2017; Shin et al., 2017; Yang and Patel, 2017; Jiang et al., 2019; Liu et al., 2019). A combination of Acrs that act via different mechanisms may be more effective at inhibiting CRISPR-Cas activity than a combination of Acrs that act redundantly (Borges et al., 2017). Thus, AcrIIA11’s novel mode of antagonism could enable more potent SpyCas9 inhibition than can be achieved using only dsDNA mimics.

Excitingly, AcrIIA11 represents just the tip of the iceberg. We have identified 50 additional metagenomic clones that putatively antagonize SpyCas9 (with nine confirmed for activity); many of these sequences are likely to contain new SpyCas9 inhibitors, *i.e.* ones not homologous to any known Acr. Functional metagenomics not only offers a powerful means to identify new *acr* genes, but also enables discovery of fundamentally new ways to inhibit Cas9. More broadly, our functional metagenomic approach allows us to detect *acr*s beyond curated sequence databases and can enable the discovery of Acrs against any CRISPR-Cas system of interest, in any microbial habitat, strain collection, or phage bank. This information will improve our understanding of phage and MGE dynamics in microbial ecosystems and holds significant promise for not only describing the longstanding evolutionary arms race between phages and bacteria, but also for improving phage- and CRISPR-Cas-based technologies used in therapeutic and engineering applications.

## Materials and Methods

### Processing of metagenomic libraries

For each metagenomic library (table S2, given generously by Gautam Dantas, Washington Univ. in St. Louis), the original freezer stock (Clemente et al., 2015; Pehrsson et al., 2016) was inoculated into 52 ml and grown to an OD600 value of ~0.7 (300 µl/100 µl inoculum for oral/fecal libraries). All libraries were previously constructed in a pZE21 MCS1 plasmid backbone (henceforth, pZE21) marked by kanamycin resistance (Kan^R^) (Lutz and Bujard, 1997; Clemente et al., 2015; Pehrsson et al., 2016). Each library was then titered on agar plates containing lysogeny broth (LB; 10 g/L casein peptone, 10 g/L NaCl, 5 g/L ultra-filtered yeast powder) with 50 ug/ml Kanamycin and 10-12 ml was used to create replicate freezer stocks. The remaining cells were pelleted by centrifugation at 4100 rcf for 6.5 minutes, purified using Qiagen miniprep kits (four columns per library), and quantified using the Qubit BR dsDNA quantification kit. The libraries Oral_3 and Oral_5 are combinations of two smaller libraries (Clemente et al., 2015); each smaller library was processed independently and then combined in proportion to the unique clones in each library before being transformed into *E. coli*.

### Cloning Cas9 expression constructs

A series of slightly different plasmids were used to express *S. pyogenes* Cas9 during the development of the Acr selection scheme and follow-up work (generically, pCas9). Plasmid composition, applications, and resultant data are detailed in tables S1, S5, and figure S2. Early versions of pCas9 were built by targeting a previously described Cas9 expression vector (addgene #48645) (Esvelt et al., 2013) to pZE21 with new crRNAs. The targeting crRNAs were engineered into the original locus (Esvelt et al., 2013) by Gibson cloning with long-tailed primers. All Gibson assembly was performed using the NEBuilder HiFi DNA assembly mastermix (NEB cat# E2621) per manufacturer’s recommendations. The arabinose-inducible *araC* and pBad regulon was amplified from pU2 (Lee et al., 2015) and, using Gibson assembly, inserted into pCas9 in place of the constitutive proC promoter to regulate Cas9 expression. To create the first 2-crRNA pCas9 construct, a gBlock was ordered from Integrated DNA Technologies (IDT) containing an I-SceI cut site and second crRNA (sequence crRNA ‘Z’ in table S5) under the control of pJ107106 and a T7TE-LuxIA terminator. This gBlock was cloned into pCas9 by Gibson assembly. Swapping crRNAs at this locus was also performed by Gibson cloning with long-tailed primers. The pJ107106 promoter was converted to pJ107111 (Zucca et al., 2015) using the Q5 site-directed mutagenesis kit (NEB), with suggested protocols. All pCas9 constructs were transformed into *Escherichia coli* (NEB Turbo) by heat-shock and made electrocompetent as described (protocols.io, 2015).

### Metagenomic selection for Cas9 antagonists

In the first iteration of Cas9 selection, 360 ng of each metagenomic library was electroporated into a strain of ultra-competent *E. coli* (NEB Turbo) containing the spectinomycin-marked Cas9 expression plasmid (pCas9). As a control, 360 ng of pZE21 expressing GFP was also transformed. All electroporations occurred in a 1 mM cuvette using a Bio-Rad Gene Pulser Xcell with the following settings: 2.1kV, 100 Ω, 25µF. Following electroporation, cells recovered for three hours at 37°C in SOC media (NEB, cat. # B9020S). An aliquot was then taken to titer transformants and the remaining cells (860 µl) were inoculated into 25 ml of LB broth with 50 ug/ml spectinomycin (Spec) and 2 mg/ml arabinose. The GFP control was split across two 25 ml flasks, one with and one without arabinose (430 µl inoculum per flask). After 20 hours in selective conditions, all samples were titered on LB-Spec or LB-Kan/Spec plates. Figure 1D depicts the population proportion of Kan^R^ colony forming units (cfu) at 20 hours after Cas9 induction relative to their proportion before Cas9 induction.

To isolate plasmids surviving one round of selection, colonies from LB-Kan/Spec titer plates (table S2) were collected in 3 ml of LB-Spec after scraping colonies with an L-shaped cell scraper (Fisher Scientific cat. # 03-392-151) to gently remove them from the agar. Colonies were collected from all metagenomic libraries and the GFP control exposed to arabinose. One-third of the cells were used to make −80°C freezer stocks and plasmids from the remaining 2 ml were purified across two Qiagen miniprep columns into a combined 100 µl of nuclease-free H_2_O. An I-SceI restriction site was engineered into pCas9 to enable its removal via treatment with the homing endonuclease. For all samples, 51 µl of miniprep eluate was combined with 6 µl NEB cutsmart buffer and 3 µl (15 units) I-SceI, incubated for 20 hours at 37°C, and the reaction heat-killed at 65°C for 20 minutes. After withholding 5 µl of each sample for gel electrophoresis, 2.98 µl of the *E. coli* RecBCD enzyme (149 units, given generously by Andrew Taylor), 4.9 µl 10mM ATP, and 1.62 µl NEB cutsmart buffer were added to the remaining 55 µl of each sample and incubated for one hour at 37°C. To kill the RecBCD reaction, EDTA was added to a final concentration of 20mM and the sample incubated at 70°C for 30 minutes. Linearization of pCas9 by I-SceI and the subsequent removal of linear DNA by RecBCD was confirmed by visualization with gel electrophoresis. Plasmid preparations were then purified through a Zymo Research DNA Clean & Concentrator column and once-selected metagenomic DNA libraries electroporated into the original Cas9 selection strain a second time (table S2). For the second iteration of Cas9 selection, electroporation, recovery, induction, titering, and outgrowth was performed exactly as described for the first iteration.

### Purification and amplification of metagenomic DNA from Cas9 selections

From twice-selected metagenomic libraries, LB-Kan/Spec titer plates were used to collect Kan^R^ clones from the population. Two titer plates were processed independently for each library and were selected based on colony counts between (roughly) 10^2^ and 10^4^ (table S2). Only one titer plate was collected for the GFP control. For each plate, colonies were collected, stored, and plasmid purified as described above. No enzymatic removal of pCas9 was performed. Instead, purified plasmids were diluted ten-fold and metagenomic DNA fragments amplified via PCR using 45µl Platinum HiFi polymerase mix (Thermo Fischer cat. # 12532016), 1.67 µl plasmid template, and 5 µl of a custom primer mix. The custom primer mix contained three forward and three reverse primers, each targeting the sequence immediately adjacent the metagenomic clone site in pZE21, staggered by one base pair. The stagger enabled diverse nucleotide composition during early Illumina sequencing cycles. A sample PCR contained the following primer volumes, each from a 10µM stock: (primer F1, 5′-CCGAATTCATTAAAGAGGAGAAAG, 0.83µl); (primer F2, 5′-CGAATTCATTAAAGAGGAGAAAGG, 0.83µl); (primer F3, 5′-GAATTCATTAAAGAGGAGAAAGGTAC, 0.83µl); (primer R1, 5′-GATATCAAGCTTATCGATACCGTC, 0.36µl); (primer R2, 5′-CGATATCAAGCTTATCGATACCG, 0.71µl); (primer R3, 5′-TCGATATCAAGCTTATCGATACC, 1.43µl). PCRs were then executed using the following conditions: 94°C for 2 minutes, 30 cycles of: 94°C for 10 seconds + 55°C for 30 seconds + 68°C for 5.5min, and 68°C for 10 minutes. The amplified metagenomic fragments were cleaned using a Zymo Research DNA Clean & Concentrator column and eluted in 80 µl of Qiagen elution buffer.

### Illumina sample preparation and sequencing

For each sample, 50 µl of PCR amplicons were diluted to a final volume of 130 µl in Qiagen elution buffer and sheared to a fragment size of approximately 500 bp using a Covaris LE220 sonicator. Sheared DNA was purified and concentrated using a Zymo Research DNA Clean & Concentrator column and eluted in 25 µl nuclease-free H_2_O. DNA was then end-repaired for 30 minutes at 25°C using 250 ng sample in a 25 µl reaction with 1.5 units T4 Polymerase (NEB cat. # M0203), 5 units T4 polynucleotide kinase (NEB cat. # M0201), 2.5 units Taq DNA polymerase (NEB cat. # M0267), 40 µM dNTPs, and 1x T4 DNA ligase buffer with 10 mM ATP (NEB cat. # B0202). Reactions were then heat-killed at 75°C for 20 minutes and 2.5 µl of 1 µM pre-annealed, barcoded sequencing adapters added with 0.8 µl of NEB T4 DNA ligase (cat. # M0202T). This reaction was incubated at 16°C for 40 minutes and heat-killed at 65°C for 10 minutes. The barcoded adapters contained a 7bp sequence specific to each sample, which allowed mixed-sample sequencing runs to be de-multiplexed. Forward and reverse sequencing adapters were stored at 1 µM in TES buffer (10mM Tris, 1mM EDTA, 50 mM NaCl, pH 8.25) and annealed by heating to 95°C and cooling at a rate of 0.1°C/sec to a final temperature of 4°C. Adapters were stored at −20°C and thawed slowly on ice before use. After adapter-ligation, samples were purified through a Zymo Research DNA Clean & Concentrator column and eluted in 12 µl of the supplied elution buffer.

Next, 6 µl from each sample was pooled together and the mixture size-selected to ~500-900 bp using a 1.9% agarose gel visualized with SYBR-Safe dye (Thermo Fisher cat. # S33102). DNA was purified from gel slices using a Zymo Research gel DNA recovery kit, eluting in 28µl nuclease-free H_2_O. Purified DNA was then enriched by PCR; a sample 25 µl reaction (in 1x Phusion HF buffer) contained 6 µl purified DNA template, 0.5 µl dNTPs (10mM), 0.25 µl Phusion polymerase (2 units/µl, Thermo Fisher cat. # F530), and 1.25 µl each of the following primers (10 µM stock): (primer F, 5’ – AATGATACGGCGACCACCGAGATCTACACTCTTTCCCTACACGACGCTCTTCCGATCT); (primer R, 5’ – CAAGCAGAAGACGGCATACGAGATCGGTCTCGGCATTCCTGCTGAACCGCTCTTCCGATCT). PCRs were amplified for 30 seconds at 98°C, subjected to 18 cycles of 98°C for 10 seconds, 65°C for 30 seconds, and 72°C for 30 seconds, and then incubated at 72°C for 5 minutes. Following enrichment PCR, samples were size-selected a second time to ~500-900 bp as previously described and purified libraries quantified using the Qubit fluorimeter (HS assay). Finally, 300 cycles of paired-end sequence data were generated at the Fred Hutchinson Cancer Research Center Genomics Core using a 17 pM loading concentration on an Illumina MiSeq machine with the v3 reagent kit.

### Assembly and annotation of functional metagenomic selections

Illumina paired-end reads were binned by barcode (perfect match required) such that data from each titer plate was assembled and annotated independently. Metagenomic DNA fragments from each sample were assembled using the first 93bp of each read with PARFuMS, a tool developed specifically for assembling small-insert functional metagenomic selections and which was optimized for shorter read lengths (Forsberg et al., 2012). The PARFuMS assembler uses three iterations of Velvet (Zerbino and Birney, 2008) with variable job size, two iterations of PHRAP (de la Bastide and McCombie, 2007), and custom scripts to clean reads, remove assembly chimeras, and link contigs by coverage and shared annotation, as previously described (Forsberg et al., 2012). In sum, 206 contigs were assembled from all ten titer plates (encompassing five selections, figure S3). We predicted open reading frames (ORFs) from each contig with MetaGeneMark (Zhu et al., 2010) using default parameters. Proteins were annotated by searching their amino acid sequences against the TIGRFAMs (Haft et al., 2001) and Pfam (Bateman et al., 2000) profile HMM databases using HMMER3 (Finn et al., 2011). The highest-scoring profile was used to annotate each ORF (minimum e-value 1e^−5^); these automated annotations were curated by hand into the functional groups shown in figure S4. The NCBI taxonomies of these contigs were determined using BLASTn against the ‘nt’ database; all contigs could confidently be assigned NCBI taxa at the resolution of order (figure S4, table S4). Seven assembled contigs matched pCas9 (>99% nucleotide identity) and were removed from the dataset. We also discarded contigs covered by less than 2 reads/base-pair/million reads (RPBM) to ensure contigs assembled from background DNA (*i.e.* plasmid backbone, host genome) were not considered (the coverage maximum among pCas9-derived contigs was 1.67 RPBM, which informed our 2 RPBM cutoff). Then, contigs which could be linked to a mutation in either protospacer or protospacer adjacent motif (PAM) were eliminated (see methods section below). Note that we use the term ‘target site’ to refer to a particular protospacer/PAM sequence. Subsequently, redundant contigs assembled across both titer plates from a given library were removed with CD-HIT (Li and Godzik, 2006), using the following parameters: -c 0.95, -aS 0.95, -g 1 (the shorter contigs among clusters sharing 95% nucleotide identity over 95% the length of the shorter contig were removed). We checked for cross-contamination among the non-redundant contigs with a BLASTn search against the unfiltered contig set, detecting (and discarding) just one cross-contaminating contig (in library Fecal_01E). Manual curation of this near-final dataset prompted removal of one chimeric contig and the fusion of two incomplete assemblies. Note that the final dataset may exclude some metagenomic clones that inhibit Cas9, specifically those with mild antagonism or high fitness costs (rare among surviving colonies, with coverage < RPBM=2) and those which have acquired target site mutation yet also encode antagonism (partial antagonism may potentiate target site mutation). Indeed, one contig (O5_7) contains a mutation in the PAM of target site B yet still confers Cas9 protection when re-cloned into a wild-type vector backbone (figure 2B). On the basis of empirical antagonism, we include this contig in our final dataset (to give 51 final contigs across five selections, tables S3, S4), but exclude all untested contigs linked to target site mutations. While these exclusions may omit *bona-fide* Cas9 antagonism, their exclusion yields a dataset with higher-confidence enrichment for Cas9 antagonists. All final assembled contigs have been submitted to GenBank.

### Determining coverage of assembled contigs

Reads used as input for each assembly (i.e. those after adapter-trimming, vector-masking, and spike-in removal by PARFuMs (Forsberg et al., 2012)) were also used to calculate coverage across each contig. For each sample, these reads were mapped to assembled contigs with bowtie2 (local alignment, unpaired reads, very-fast mode) and RPBM calculated for each contig.

### Determining the proportion of wild-type target sites

To estimate the proportion of wild-type target sites in each library after two rounds of Cas9 selection, titer-specific reads (before vector-cleaning) were concatenated by library and mapped to the pZE21 vector with bowtie2 (end-to-end alignment, unpaired reads, sensitive mode). For the GFP control, reads were used from the only titer plate collected. Base frequencies at each protospacer and PAM position were calculated using bam-readcount (github.com/genome/bam-readcount), requiring minimum mapping and base quality scores of 25 and 30, respectively. The proportion of wild-type target sites was conservatively calculated as the product of wild-type base calls at each protospacer/PAM position. In two libraries, (Oral_3 and Fecal_01E), sanger sequencing of individually picked colonies revealed a recombination event between a metagenomic DNA fragment and target site B, which would remove this Cas9 target from these clones (but avoid frameshift within the Kan^R^ gene). For these two libraries, reads were mapped to a variant of the pZE21 backbone corresponding to each recombination mutant. The proportion of reads mapping across the recombination site was compared to the proportion mapping to the wild-type vector backbone to estimate the frequency of recombination mutants in the population (0.4% and 24.2% for Oral_3 and Fecal_01E, respectively). The overall proportion of wild-type target sites in these libraries was then adjusted to reflect the proportion of these recombinants (table S3).

### Linking contigs to target site genotype

For some of the most abundant contigs in the final dataset, the genotype of the target site sequences in the corresponding plasmid backbone could be determined by Sanger sequencing individual Kan^R^ clones following two rounds of Cas9 selection. To link additional contigs with target site genotype in the clones not sampled by our Sanger data, we examined paired-end reads that mapped to target site sequences and an assembled contig. Because we used PCR to amplify metagenomic fragments before Illumina library creation, only a small minority of reads mapped to the vector backbone. Additionally, target sites A and B are 514 bp and 365 bp from the clone site of pZE21, respectively, so only read pairs from long sequencing inserts could link target site and contig genotype. Accordingly, many assembled contigs remain without links to target site genotype (figure S3), though the overall proportion of wild-type target sites serves as a proxy for the number of Cas9 antagonists versus escape mutants in a given library (table S3).

For each titer plate, paired-end reads were mapped to a reference database containing all assembled contigs in both forward and reverse orientations within the pZE21 plasmid. We used bowtie2 in “no-mixed” mode to consider only paired reads with an insert size ≥515 bp when mapping against this reference database (additional mapping options: end-to-end alignment, paired reads, sensitive mode). Only reads with a mapping quality score over ten were considered. We determined the true orientation of a metagenomic DNA fragment by counting whether more reads mapped across the clone-site junctions of the contig in the forward or reverse orientation. If more than one mapped read supported a particular protospacer/PAM variant, the contig linked to that target site was removed from the final dataset. Similarly, if fewer than ten reads mapped to a target site, a single read with any deviation from a wild-type protospacer/PAM was sufficient to eliminate the corresponding contig from further consideration.

### Predicting phage-associated contigs

The assembled metagenomic contigs (N50 length 2.2 Kb) were too short to provide sufficient information content for phage prediction based on their nucleotide sequence (Roux et al., 2015). Instead, protein sequences were used: those with close relatives in predicted phages were used to classify their parent contig as phage-associated. Predicted proteins from each contig were queried against NCBI’s NR database using BLASTp (on September 12^th^, 2017) and the top five unique hits retained from each query. For each BLAST hit, we downloaded sequence 25 Kb upstream and downstream of the corresponding gene sequence and used these ~50KB sequences as input for phage prediction with VirSorter (Roux et al., 2015). Sequences were compared against VirSorter’s more encompassing ‘virome’ database (using default parameters) and hits to bacteriophage of any confidence category (from ‘possible’ to ‘most confident’) were considered phage-associated (Roux et al., 2015). The proteins that seeded these bacteriophage predictions were used then to classify metagenomic DNA contigs. If any predicted protein from our assembled contigs had a top-five BLAST hit that seeded a bacteriophage prediction, the whole metagenomic contig was classified as being phage-associated. These classifications are referenced in figure 2A and table S4.

### Validating contigs with Cas9 protection

All contigs assembled from libraries Fecal_01A and Oral_5 with coverage above the sample median (RPBM >3.6, Fecal_01A and RPBM > 8.2, Oral_5) were chosen for validation, totaling 15 contigs. Five contigs were represented in a sparsely-sampled set of individually-picked and Sanger-sequenced colonies. In these cases, the corresponding plasmids were re-transformed into NEB Turbo, target site sequences confirmed as wild-type by additional Sanger Sequencing, and the plasmids co-transformed with pCas9 into NEB Turbo for functional testing. The remaining contigs were amplified by PCR from the plasmid preparations that followed two iterations of SpyCas9 selection (those used in PCRs for Illumina library construction). PCR primers (table S6) targeting the junction between pZE21 and the assembled contigs were used to facilitate re-cloning of the amplified fragments into pZE21 by Gibson assembly. PCRs used the high-fidelity Q5 polymerase (NEB) per suggested protocols. Cycling conditions are as follows: 98°C for 3 minutes, 35 cycles of 98°C for 15 seconds + 61°C or 65°C for 30 seconds + 72°C for 3 minutes, and 72°C for 10 minutes. Reaction-specific annealing temperatures are listed in table S6 and PCR products of the expected size were purified from gel slices using a Zymo Research gel DNA recovery kit. Nine of ten PCRs were successful [a contig from Fecal_01A (RPBM=3.66) did not amplify well]. All Gibson assembly was performed using the NEBuilder HiFi DNA assembly mastermix per manufacturer’s recommendations. Following Gibson-assembly, sequence-verified clones were co-transformed with pCas9 into NEB Turbo, resulting in 14 total strains re-tested for Cas9 antagonism.

These 14 strains and a control carrying an empty pZE21 vector were grown overnight in LB with 50 µg/ml kanamycin and spectinomycin (LB-Kan/Spec) to maintain the pZE21 variants and pCas9, respectively. The next morning, cultures were diluted 30-fold into LB-Kan/Spec and grown to log phase (absorbance readings at 600 nm ranged from 0.2 to 0.4). Mid-log cultures were then inoculated into LB with 50 ug/ml spectinomycin and either 0.2 mg/ml arabinose (to induce Cas9) or no arabinose. Kanamycin was omitted to allow for restriction of the pZE21 plasmid. Inocula were normalized by optical density (a 1:40 inoculum was used for an absorbance reading of 0.4). Cultures were then arrayed in triplicate across a 96-well plate (Greiner cat#655083, 200µl per well) and grown in a BioTek Cytation 3 plate reader at 37°C with linear shaking at 1096 cycles per minute (cpm). After six hours of growth (at which time cultures had reached stationary phase), each well was serially-diluted ten-fold in peptone-NaCl (1 g/L each, pH 7.0). Spots (5 µl) of each dilution were plated on agar plates with LB-Kan/Spec (to count Kan^R^ colonies) and LB-Spec (to count total colonies). Figure 2B depicts the log-transformed proportion of Kan^R^/total cfu with and without Cas9 induction.

### Identification of causal Acr proteins

Null alleles of each predicted ORF in F01A_2 were created by inserting an early stop codon into each predicted protein. Stop codons were inserted by PCR using the high-fidelity Q5 polymerase (NEB) per suggested protocols with the following cycling conditions: 98°C for 3 minutes, 35 cycles of 98°C for 15 seconds + [annealing temperature] for 30 seconds + 72°C for [extension time], and 72°C for 10 minutes. Reaction-specific primers and conditions are listed in table S6. Constructs were then prepared using the Q5 site-directed mutagenesis kit (NEB cat #E0554) or by Gibson assembly (using the NEBuilder HiFi assembly mastermix, cat# E2621) per manufacturer’s recommendations. Final constructs were sequence-verified, co-transformed with pCas9 into NEB Turbo, and re-tested for Cas9 antagonism alongside an empty-vector control and the parent pZE21-F01A_2. Plasmid protection assays were performed exactly as described in the section titled ‘Validating contigs with Cas9 protection’, except that Cas9 expression was induced using 2 mg/ml arabinose (rather than 0.2 mg/ml).

### Cloning candidate Acrs

The third ORF (*acrIIA11*) from F01A_2 and *acrIIA4* were codon-optimized for *E. coli* and sub-cloned into the KpnI (5’) and HindIII (3’) sites of a pZE21 containing the *tetR* gene to allow for doxycycline-induced expression of the candidate Acrs from the pLtetO-1 promoter. To generate this plasmid, *tetR* (and its pLac promoter + rrnB terminator) was amplified from pCRT7 (addgene #52053) with NEB’s Q5 polymerase per suggested protocols and using the conditions listed in table S6. This amplicon was then cloned by Gibson assembly into the pZE21 backbone. To clone candidate Acrs, KpnI and HindIII sites were added to each gene by PCR with the Q5 polymerase (see table S6), vectors and genes digested using these enzymes, and sticky-end ligations performed using the Fast-Link ligation kit (epicentre cat# LK0750H) per suggested protocols. The *acrIIA11 and acrIIA4* gene sequences were synthesized as gBlocks from IDT (the AcrIIA4 amino acid sequence is identical to NCBI accession AEO04689.1). AcrIIA11 homologs were codon-optimized for *E. coli*, synthesized by GenScript, and cloned into this identical vector context. All constructs were co-transformed with pCas9 into NEB Turbo prior to performing plasmid protection assays. Because NEB Turbo contains the *laci^q^* mutation, 0.5 mM IPTG was used throughout cloning and handling of the pZE21_tetR plasmid in this strain. IPTG was used because *TetR* is driven by the pLac promoter in this construct. Including IPTG ensured that TetR was at sufficient intracellular concentrations to maintain repressed transcription from the pLtetO-1 promoter prior to Acr induction.

### Testing candidate Acrs for plasmid protection

Overnight cultures of candidate Acr constructs were grown in LB-Kan/Spec + 0.5mM IPTG (LB-Kan/Spec/IPTG) to maintain pCas9 and the Acr variants in an uninduced state. The next morning, cultures were diluted 1:50 into LB-Kan/Spec/IPTG and grown at 37°C for approximately two hours to mid-log with or without 100 ng/ml doxycycline to control Acr expression (final absorbance readings ranged from 0.2 to 0.6). Mid-log cultures were then inoculated into LB with 50 ug/ml spectinomycin, 0.5 mM IPTG, and either 2 mg/ml arabinose (to induce Cas9) or no arabinose. Doxycycline was included in this medium at 100 ng/ml to maintain Acr expression only for previously induced cultures. Kanamycin was omitted to allow for elimination of the pZE21 target plasmid and inocula were normalized by optical density (a 1:40 inoculum was used for an absorbance reading of 0.4). Cultures were grown in a 96-well plate reader and the proportion of Kan^R^ cfu determined exactly as described in the section ‘Validating contigs with Cas9 protection’. Figures 3 and 4 depicts the log-transformed proportion of Kan^R^/total cfu with and without Cas9 induction for each Acr.

### Phage plaquing assay

Overnight cultures with pCas9 and pZE21_tetR expressing either GFP or an Acr were diluted 1:50 in LB-Kan/Spec/IPTG supplemented with 5 mM MgSO_4_ and grown shaking at 37C for three hours until late-log phase (OD600 range 0.5-0.8, see table S5 for exact crRNA sequences). GFP and the candidate Acrs were induced by adding doxycycline to a final concentration of 100 ng/ul and cultures grown for another two hours before SpyCas9 was induced by adding 0.2 mg/ml arabinose. After three additional hours of growth in the presence of both inducers, 200µl of culture was used in a top-agar overlay, allowed to harden, and ten-fold serial dilutions of phage Mu spotted on top. The top and bottom agar media were made with LB-Kan/Spec/IPTG supplemented with 5mM MgSO4, 0.02 mg/ml arabinose, and 100 ng/ul doxycycline and contained 0.5% and 1% Difco agar, respectively. Plates were incubated at 37°C overnight and plaques imaged the following day.

### AcrIIA11 homolog discovery and phylogenetic placement

To determine the distribution AcrIIA1, AcrIIA2, AcrIIA3, AcrIIA4, AcrIIA5, AcrIIA6, and AcrIIA11 across bacteria (figures 3 and S7), homologs of each Acr were identified via a BLASTp search against NCBI nr; all sequences with ≥35% amino acid identity over ≥75% of the query length were retrieved. Many AcrIIA11 homologs were derived from bacteria with phylogenetically-ambiguous taxonomic labels in NCBI. For instance, the NCBI genus ‘Clostridium’ is extremely polyphyletic, appearing in 121 genera and 29 families in a rigorous re-assessment of bacterial phylogeny (work via the genome taxonomy database, or GTDB (Parks et al., 2018)). Importantly, the GTDB taxonomy ensures that taxonomic labels are linked to monophyletic groups, applies taxonomic ranks (phylum, class, order, etc.) at even phylogenetic depths, and substantially improves problematic taxonomies (Parks et al., 2018). To make phylogenetically meaningful inference with respect to AcrIIA11, we adopted the GTDB taxonomic scheme which, crucially, is largely congruent with NCBI taxonomy outside a few key problem areas (e.g. Clostridiales, see table S4). The taxonomic assignments for AcrIIA1-6 homologs are unchanged between NCBI and GTDB schema.

To map protein homologs onto a bacterial genus tree, the NCBI genome assemblies matching each homolog were downloaded and classified according to the GTDB scheme using the ‘classify_wf’ workflow within GTDB toolkit (v0.1.3) (Parks et al., 2018); default parameters were used. To visualize these classifications on a tree of life, the minimal phylogenetic tree encompassing all lineages encoding AcrIIA1-6 or AcrIIA11 homologs was downloaded from AnnoTree (v1.0; node ID 31285). The AnnoTree phylogeny additionally includes KEGG and PFAM annotations for nearly 24,000 bacterial genome assemblies and is classified according to GTDB taxonomy (Mendler et al., 2018). Node 31285 includes five bacterial phyla which are identified with letters on figure S7 (a: Firmicutes, b: Firmicutes_D, c: Firmicutes_A, d: Fusobacteria, and e: Firmicutes_F). The KEGG identifier K09952 was used to determine which GTDB genera encoded Cas9 while the PFAM IDs PF09711 and PF16813 were used to identify taxa which encode Csn2, the signature protein of type II-A CRISPR-Cas systems (Makarova et al., 2015). Enrichment for type II-A CRISPR-Cas systems was determined via a chi-squared test using the number of Csn2-encoding genomes within each GTDB family, with p-values corrected for multiple hypotheses using the Bonferroni correction.

### Phylogenetic tree of AcrIIA11 homologs

The gene tree in figure S4 was built using the homologs in NCBI (see above section) and additionally a set of homologs identified via a BLASTP search of IMG/VR (the January 1, 2018 release), a curated database of cultured and uncultured DNA viruses (Paez-Espino et al., 2017). An e-value cutoff of 1×10^−10^ was used in the IMG/VR homolog search. A preliminary phylogeny of all AcrIIA11 homologs was used to select genes that optimally sample AcrIIA11 diversity for gene synthesis and anti-SpyCas9 activity screening. The final phylogeny in figure S4 contains all NCBI homologs and the viral homologs which were selected for gene synthesis (see table S7 for sequences and accession numbers). Alignments were performed using the Geneious (v8) alignment tool and a maximum-likelihood tree was generated with PhyML using the LG substitution model and 100 bootstraps.

### Protein expression and purification

Codon-optimized AcrIIA4 and AcrIIA11 were cloned into pET15b to contain thrombin-cleavable 6XHis N-terminal tags, with and without C-terminal 2xStrep2 tags. All plasmids were transformed into *E. coli* BL21(DE3) RIL cells except the C-terminally-tagged AcrIIA11 variant, which was transformed into a BL21(DE3) pLysS strain. Overnight cultures were grown in LB/Ampicillin (100 µg/mL), diluted 100-fold into 1 L pre-warmed LB/Amp media, grown until an OD600 of 0.6-0.8, then incubated on ice for 30 minutes. IPTG was added to 0.2 mM, and the cultures where shaken for 18-20 hours at 18°C. The cells were pelleted and stored at −20°C. Cell pellets were resuspended in lysis/wash buffer (500 mM NaCl, 25 mM Tris, pH 7.5, 20 mM Imidazole), lysed by sonication, and centrifuged for 25 minutes in an SS34 rotor at 18,000 rpm for 30 minutes. The soluble fraction was filtered through a 5 µm filter and incubated in batch with Ni-NTA resin (Invitrogen, Cat# R90115) at 4°C for 1 hour. The resin was transferred to a gravity filtration column and washed with at least 50 volumes of wash buffer, followed by elution in 200 mM NaCl, 25 mM Tris, pH 7.5, 200 mM Imidazole. The buffer was exchanged into 200 mM NaCl/25 mM Tris, pH 7.5 by concentrating and diluting using an Amicon filter (EMD Millipore, 10,000 MWCO). Biotinylated thrombin (EMD Millipore) was added (1 U per mg of protein) and incubated for 16 hours at 18°C. Streptavidin-agarose was then used to remove thrombin according to the manufacturer’s instructions (EMD Millipore). The proteins were diluted to 150 mM NaCl, 25 mM Tris pH 7.5, loaded onto a 1 mL HiTrapQ column (GE Life Sciences), and eluted by a sodium chloride gradient (150 mM to 1 M over 20 mL). Peak fractions were pooled and concentrated, bound to Ni-NTA to remove any uncleaved proteins, and the flow-through was purified via size exclusion chromatography (using Superdex75 16/60 (GE HealthCare) or SEC650 (BioRad) columns) in 200 mM NaCl, 25 mM Tris, pH 7.5, 5% glycerol. Figure 5A depicts size exclusion chromatography data for thrombin-cleaved AcrIIA11 lacking a 2xStrep2 tag. Peak fractions were pooled, concentrated, flash frozen as single-use aliquots in liquid nitrogen, and stored at −80°C.

Plasmid pMJ806 (addgene #39312) was used to express SpyCas9 containing an N-terminal 6XHis-MBP tag followed by a TEV protease cleavage site. The protein was expressed and purified over Ni-NTA as described above. The eluate from the Ni-NTA resin was dialyzed into 25 mM Tris, pH 7.5, with 300 mM NaCl, 1 mM DTT, and 5% glycerol. It was simultaneously cleaved overnight with homemade TEV protease at 4°C. The cleaved protein was loaded onto a 1 mL heparin HiTrap column (GE) in 300 mM NaCl, 25 mM Tris, 7.5 and eluted in a gradient extending to 1 M NaCl/ 25 mM Tris, 7.5. Pooled fractions were concentrated and buffer-exchanged into with 200 mM NaCl, 25 mM Tris (pH 7.5), and 20 mM imidazole, and bound to Ni-NTA to remove uncleaved fusion proteins. The flow-through was purified over a Superdex 200 16/60 column (GE Healthcare) equilibrated in buffer containing 200 mM NaCl, 25 mM Tris (pH 7.5), 5% glycerol, and 2 mM DTT. Peak fractions were pooled, concentrated to 3-6 mg/mL, flash frozen as single-use aliquots in liquid nitrogen, and stored at −80°C. A second SpyCas9 variant was also expressed and purified from a pET28a backbone (addgene #53261) as described above except that no protease cleavage site was present, so tags were not removed. This second SpyCas9 variant includes an N-terminal 6xHis tag, a C-terminal HA-tag, and a C-terminal NLS sequence. DNA cleavage assays and Acr pulldowns were performed using tagged SpyCas9, whereas EMSAs used the untagged version. EMSAs were also performed using the tagged SpyCas9 variant and the results did not differ between SpyCas9 purifications.

### gRNA Generation

gRNA was generated by T7 RNA polymerase using Megashortscript Kit (Thermo Fisher #AM1354). Double-stranded DNA template was generated by a single round of thermal cycling (98°C for 90 seconds, 55°C for 15 seconds, 72°C for 60 seconds) in 50 µl reactions using Phusion PCR polymerase mix (NEB) containing 25 pmol each of the following ultramers (the protospacer-matching sequence is underlined): GAAATTAATACGACTCACTATAGG<u>TAATGAAATAAGATCACTAC</u>GTTTTAGAGCTAGAAATAGCAAGTTAAAATAAGGCTA GTCCG and AAAAAAGCACCGACTCGGTGCCACTTTTTCAAGTTGATAACGGACTAGCCTTATTTTAACTTGC.

The dsDNA templates were purified using an Oligo Clean and Concentrator Kit (ZymoResearch) and quantified by Nanodrop. Transcription reactions were digested with DNAse, extracted with phenol-chloroform followed by chloroform, ethanol precipitated, resuspended in RNase free water and stored at −20°C. RNA was quantified by Nanodrop and analyzed on 15% acrylamide/TBE/UREA gels.

### DNA Cleavage Assay

The buffer used in DNA cleavage reactions was NEB buffer 3.1 (100 mM NaCl, 50 mM Tris-HCl, pH 7.9, 10 mM MgCl2, 100 µg/mL BSA); proteins were diluted in 130 mM NaCl, 25 mM Tris, pH 7.4, 2.7mM KCl. SpyCas9 (0.4 µM) and AcrIIA11 (0.4 – 12.8 µM) were incubated for 10 minutes at room temperature before the reaction was started by simultaneously adding 0.4 µM gRNA and 4 nM linearized plasmid (2.6 Kb) and transferring reactions to a 37°C water bath. After 10 minutes at 37°C, the reaction was stopped by adding 0.1% SDS and 50 mM EDTA. Reactions were then run on a 1.25% agarose gel containing ethidium bromide at 115V for 2 hours at room temperature. Gels were imaged using the ethidium bromide detection protocol on a BioRad Chemidoc gel imager.

### Pull-down assays using Strep-tagged AcrIIA4 and AcrIIA11

The binding buffer for pull-down assays was 200 mM NaCl, 25 mM Tris (pH 7.5); protein dilutions were made in the same buffer. In 20 µl binding reactions, 90 pmol of SpyCas9 and gRNA were incubated for 20 minutes at room temperature, followed by incubation with 150 pmol of strep-tagged Acr for an additional 20 minutes at room temperature. 50 µl of a 10% slurry of Streptactin Resin (IBA biosciences #2-1201-002) equilibrated in binding buffer was added to the binding reactions and incubated at 4°C on a nutator. Thereafter all incubations and washes were carried out at 4°C or on ice. The beads were washed a total of four times, including one tube transfer, by centrifuging 1 minute at 2000 rpm, carefully aspirating the supernatant with a 25 gauge needle and resuspending the beads in 100 µl binding buffer. After the final bead aspiration, Strep-tagged proteins were eluted by resuspending in 40 µl of 1X BXT buffer (100 mM Tris-Cl, 150 mM NaCl, 1 mM EDTA, 50 mM Biotin, pH 8.0) and incubated for 15 minutes at room temperature. The beads were spun and 30 µl of the supernatant was carefully removed and mixed with 2X SDS Sample Buffer (Novex). Proteins were then separated by SDS PAGE on BOLT 4-12% gels in MES buffer (Invitrogen), followed by Coomassie staining. In instances where DNA was also visualized, the gel was imaged using the DyLight 488 detection protocol on a BioRad Chemidoc gel imager (Coomassie staining followed DNA visualization using the same gel).

### Electrophoretic Mobility Shift Assays

Reactions were carried out in EMSA binding buffer (56 mM NaCl, 10 mM Tris, pH 7.4, 1.2 mM KCl, 5% glycerol, 1 mM DTT, 2 mM EDTA, 50 µg/ml heparin, 100 µg/ml BSA, 0.01% Tween-20); proteins were diluted in 130 mM NaCl, 25 mM Tris, pH 7.4, 2.7mM KCl. Omitting MgCl2 ensured that Cas9 did not cleave target DNA, as previously described (Lee et al., 2018). DNA gel shifts used the 6FAM-labeled 60-mer target dsDNA (see table S6 for sequence) as template and was visualized on a BioRad Chemidoc gel imager as described for the pulldowns. All incubations were carried out at room temperature. Cas9 and gRNA (each at 2 µM) were incubated for 25 minutes, followed by addition of Acrs (2-16 µM) for 20 minutes, followed by addition of 20 nM dsDNA template and incubation for 20 minutes. Samples were loaded on an 8% acrylamide/0.5X TBE gel that was pre-run (30 minutes, 90 V, 4°C), and resolved for 160 minutes, 4°C, 90 V in 0.5X TBE buffer. For gRNA EMSA experiments, AcrIIA11 (at 4-32 µM) was incubated with 2 µM Cas9 for 20 minutes, followed by incubation with 0.2 µM gRNA for 20 minutes. The same buffers were used as in DNA EMSAs, except that 3 mM MgCl2 was included in the reactions. Samples were run for 180 minutes under the gel conditions described above. The gels were post-stained with a 1:10,000 dilution of SYBR-Gold (Invitrogen) in 0.5X TBE to visualize RNA.

## Acknowledgements

We thank Cara Forsberg, Tera Levin, Courtney Schroeder, Gerry Smith, Jeannette Tenthorey, and Janet Young for comments on the manuscript, Gautam Dantas for the functional metagenomic libraries, Andrew Taylor for purified RecBCD, and the Fred Hutchinson Cancer Research Center Genomics core facility for Illumina sequencing. This work was supported by a postdoctoral fellowship awarded to KJF by the Helen Hay Whitney Foundation and by a Seattle University summer faculty fellowship to BKK. Further support for this work includes discretionary funding from the Fred Hutchinson Cancer Research Center to BLS and grants from the National Institutes of Health to BLS (R01 GM105691) and from the Howard Hughes Medical Institute to HSM. The funders played no role in study design, data collection and interpretation, or the decision to publish this study. HSM is an Investigator of the Howard Hughes Medical Institute.

## Competing Interests

All authors declare no significant competing financial, professional, or personal interests that might have influenced the performance or presentation of the work described in this manuscript.

---

**Figure S1.**
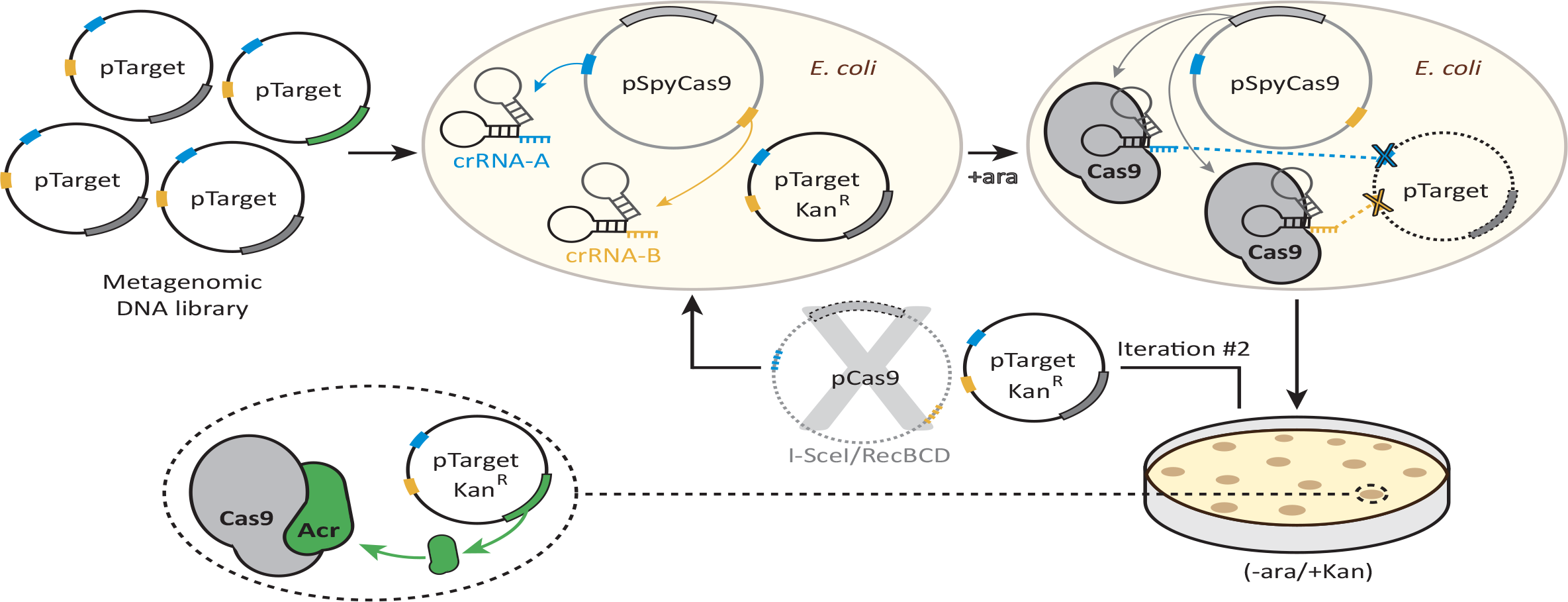
A functional selection for type II-A anti-CRISPRs. A library of plasmids each bearing a different metagenomic DNA fragment is transformed into an *E. coli* strain expressing SpyCas9 and two crRNAs, which target the library for destruction. Two crRNAs targeting two sites in the plasmid backbone reduce the number of target-site escape mutations, mitigating this source of false-positives. After transformed cells are allowed to recover, SpyCas9 is induced with arabinose and the library is subjected to SpyCas9 selection for twenty hours. Plasmids that survive this first round of selection are purified from Kan^R^ clones and the pCas9 plasmid is removed via digestion with I-SceI and RecBCD treatment. The metagenomic library is then subjected to SpyCas9 exposure a second time, which enriches for plasmid-intrinsic SpyCas9 resistance (*i.e.* what may be encoded by the metagenomic DNA inserts). The second iteration allows *acr*-encoding clones to be enriched above background, which is set by the frequency of Cas9 loss-of-function mutations. Kan^R^ clones following two rounds of selection were then harvested and their metagenomic DNA inserts sequenced to identify putative *acr*-encoding metagenomic DNA fragments.

**Figure S2.**
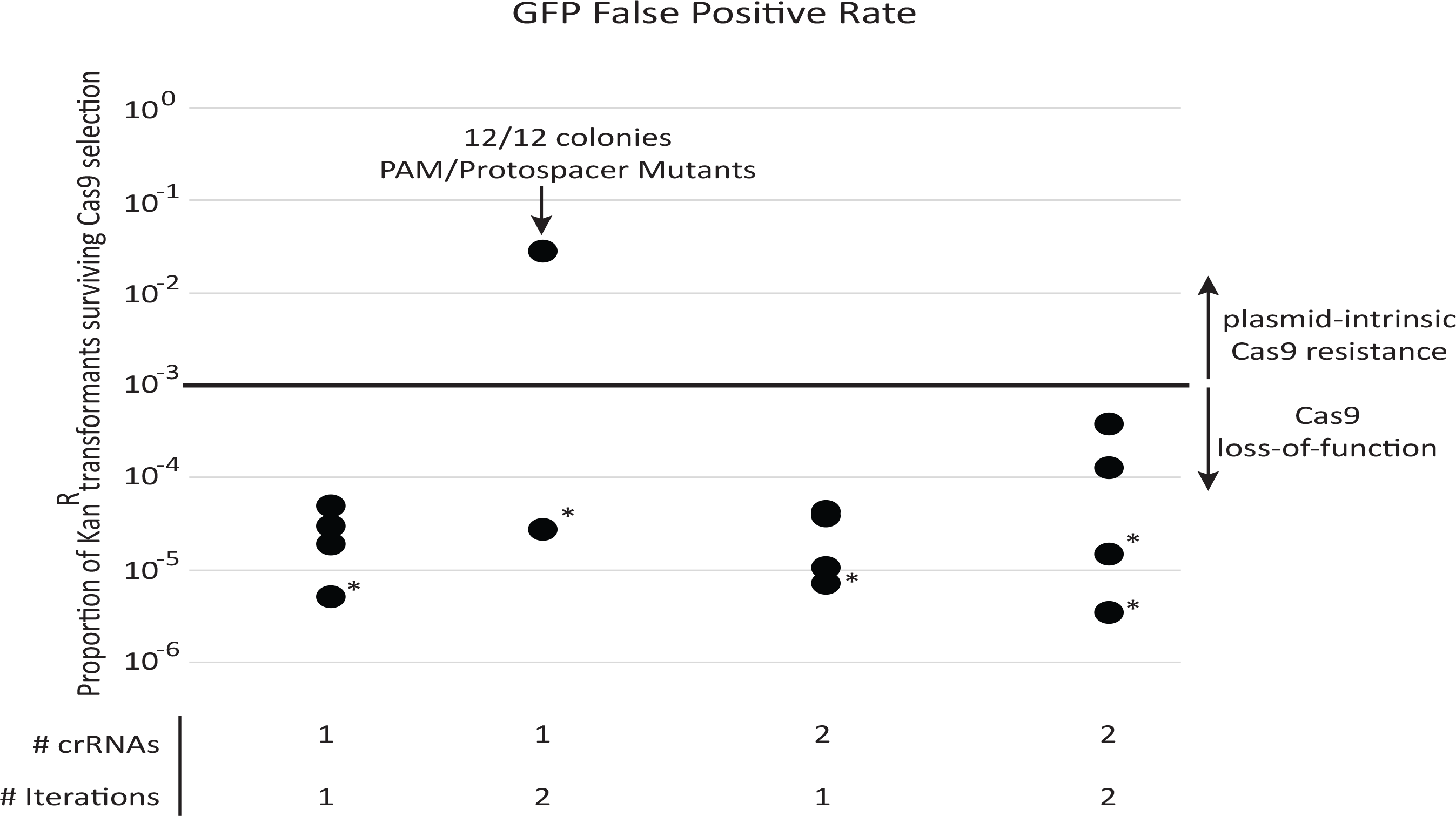
Cas9 loss-of-function mutations are the major source of false-positives during *acr* selection. Each data point represents a separate experiment toward developing the final selection for SpyCas9 antagonists. All transformations use the pZE21-GFP control target plasmid and approximate a metagenomic library expressing only neutral functions. Surviving colonies therefore represent sources of false positives. A single iteration of SpyCas9 exposure reduces Kan^R^ transformants by a factor of 10^4^ to 10^5^. This false-positive rate remained constant across all experiments except for one experiment that used a single target site on pZE21. In this experiment, mutations to the protospacer or PAM region of the Cas9 target site in pZE21 dominated, prompting two target loci to be used thereafter. All other colonies genotyped (those in asterisked experiments) escaped selection due to inactivating mutations in pCas9. This loss-of-function rate, importantly, remained constant across rounds of selection, allowing for plasmid-intrinsic Cas9 resistance to be identified via two iterations through SpyCas9 selection. When libraries were subject to two rounds of selection, this plasmid-intrinsic resistance was predominantly due to genes encoded by its metagenomic DNA fragment (fig. 1D).

**Figure S3.**
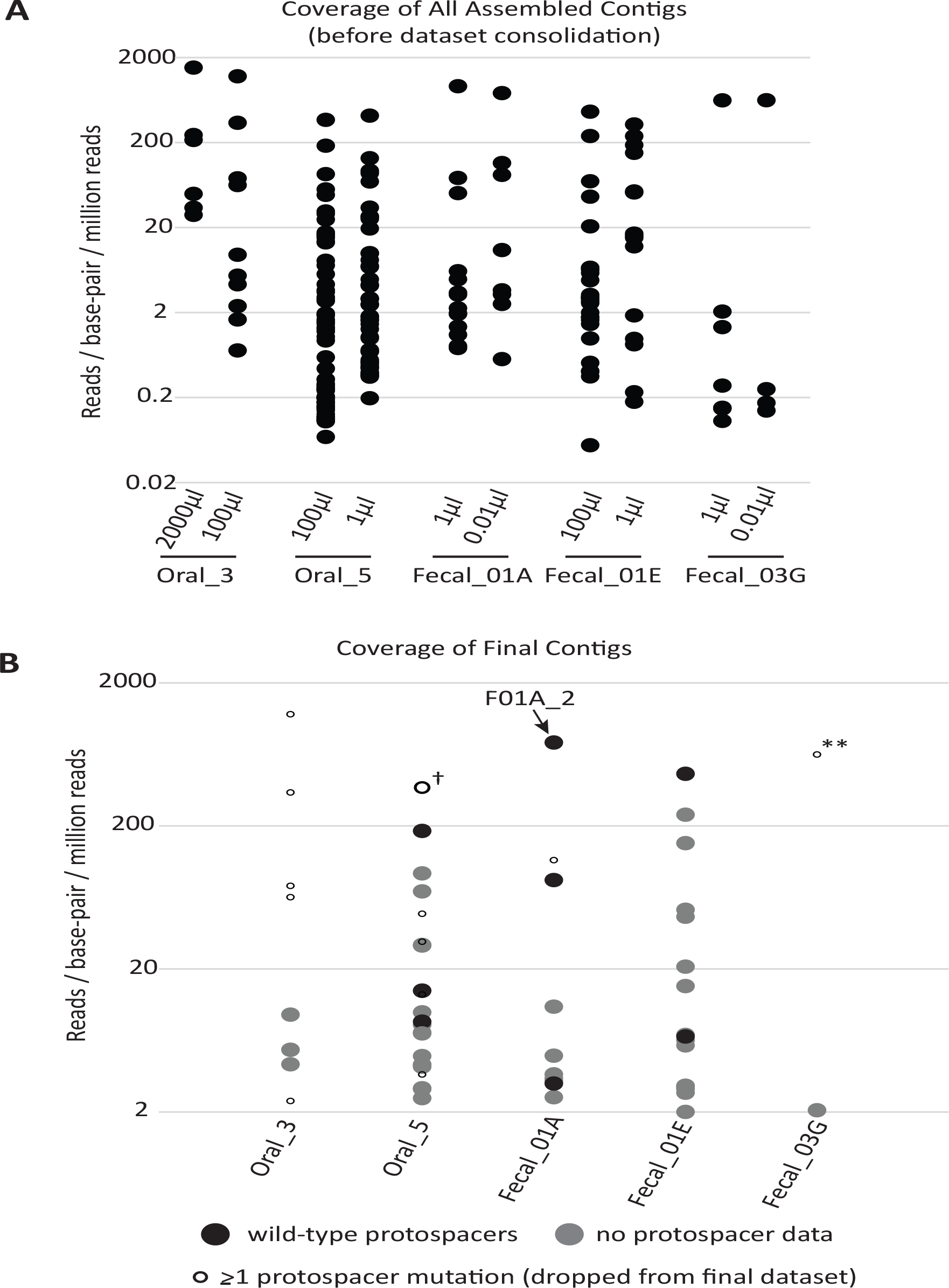
Coverage of assembled contigs by library. **(A)** Coverage of each assembled contig before dataset processing. The contigs from each titer plate were sequenced separately and assembled independently. Redundant contigs were removed during data processing to give the final contigs depicted in (B). Relative coverage can be used as a proxy for contig abundance within a library but cannot be used for meaningful comparisons across libraries. **(B)** Coverage of contigs in the final dataset. Filled circles depict contigs reported in the final dataset (n=51); grey indicates contigs without genotypic data for Cas9 target sites and black indicates contigs from plasmids with confirmed wild-type target sites. Small, empty circles depict assembled contigs linked to at least one target site mutation and were not included in the final dataset, with the cross (†) indicating the lone exception. This contig was found in a plasmid containing a single target site mutation but nonetheless was confirmed for anti-SpyCas9 activity when cloned into a fresh plasmid background, so was included in the final dataset. The double asterisks (**) highlight a clone with escape mutations in both target sites that dominated the selection using library Fecal_03G.

**Figure S4.**
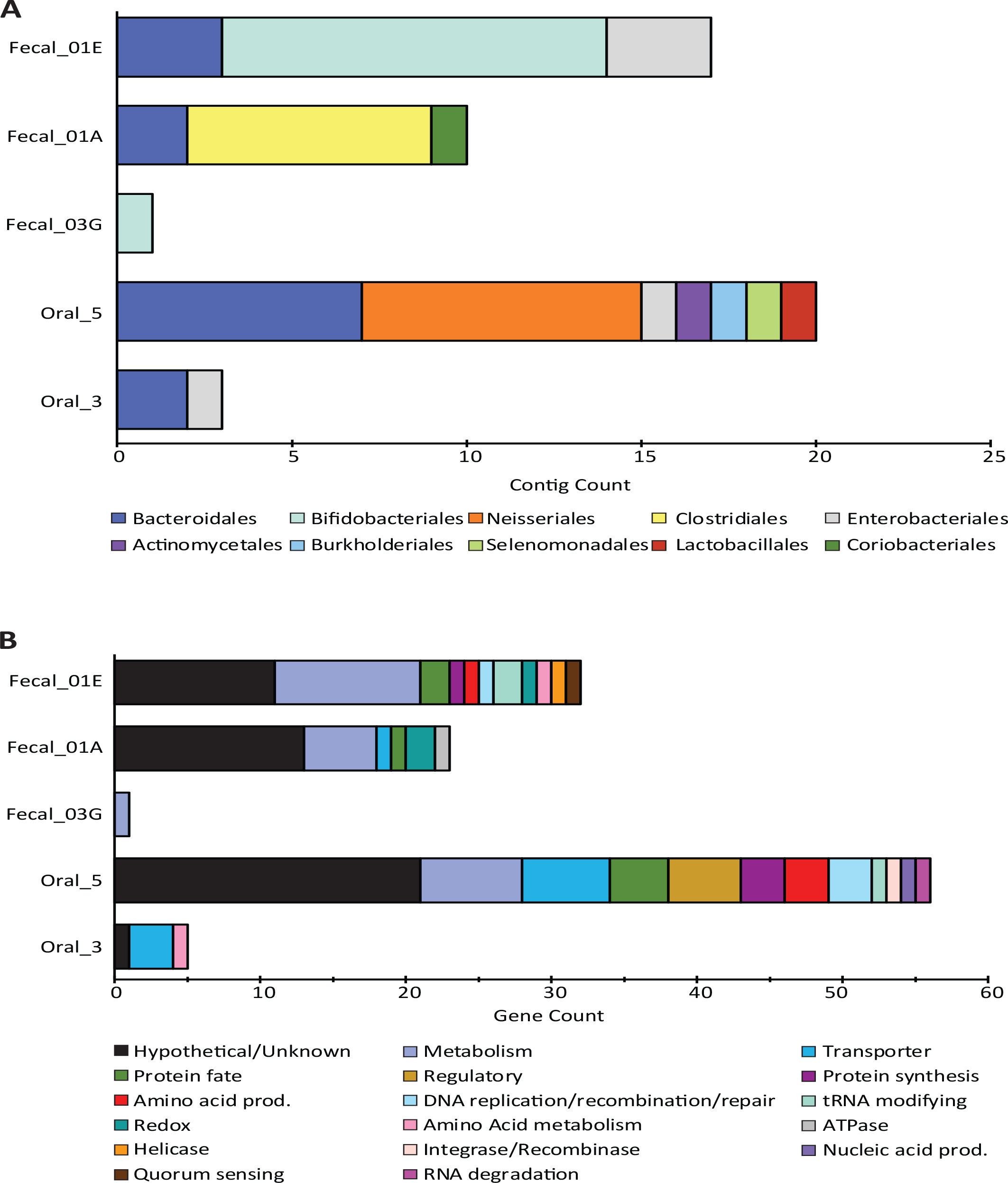
NCBI taxonomies and gene functions recovered from all SpyCas9-antagonizing contigs. **(A)** Taxonomic orders for the contigs surviving SpyCas9 targeting, grouped by library. **(B)** The encoded gene functions on these contigs, hand-curated into broad categories.

**Figure S5.**
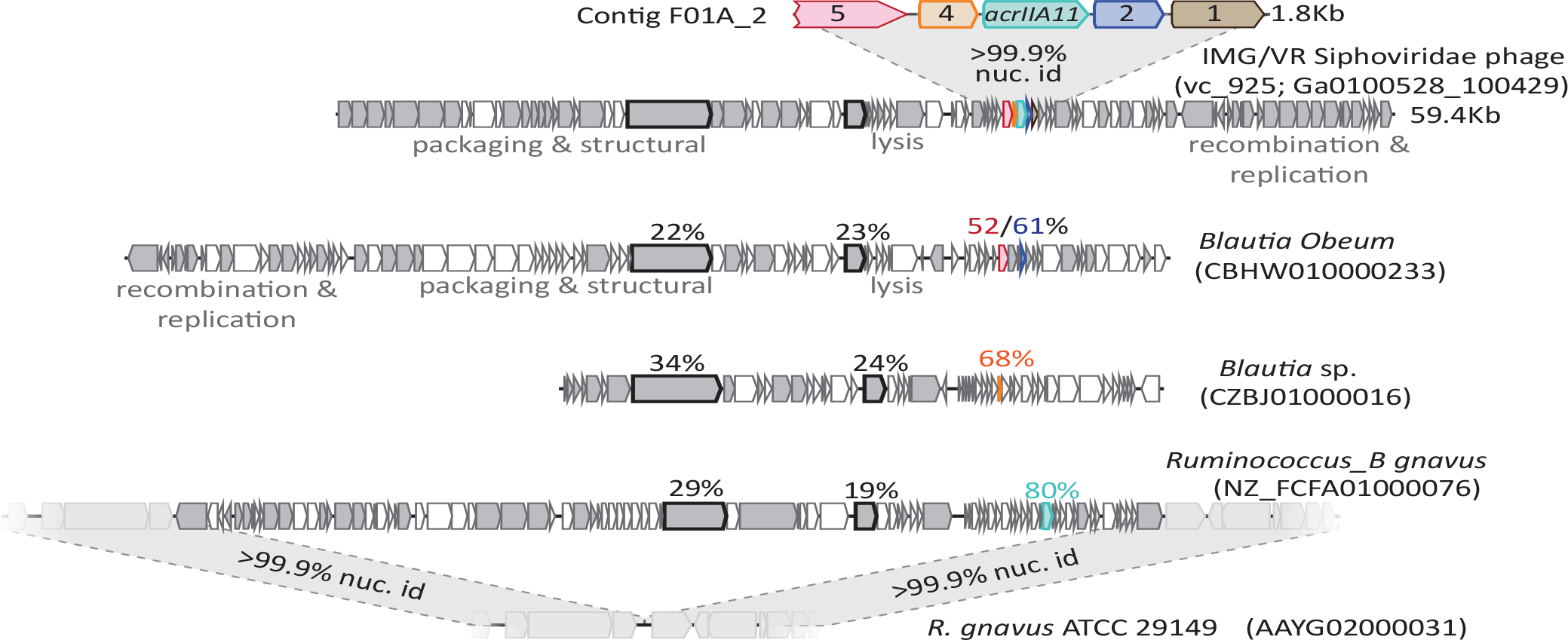
The genes on contig F01A_2 are small accessory proteins of *Lachnospiraceae* phages. These genes move readily by horizontal gene transfer and are typically unlinked from one another. Gray genes denote homologs shared between the AcrIIA11-encoding Siphoviridae phage and the three other related phages. The amino acid identities (relative to the Siphoviridae phage) for two genes present in all genomes, a tape measure protein and reverse transcriptase, are depicted in bold outline to illustrate phage relatedness. Homologs of F01A_2 genes are indicated by common color and the amino acid identities for each gene product are depicted above each gene. The depicted homolog of *acrIIA11* (named *acrIIA11a.2*) is found in an actively circulating temperate phage of *Ruminococcus gnavus*. All accession numbers denote NCBI Genbank IDs except for the Siphoviridae phage (for this phage, we use IMG/VR convention and indicate viral_cluster; scaffold_id in parentheses). Contig lengths are depicted next to each sequence.

**Figure S6.**
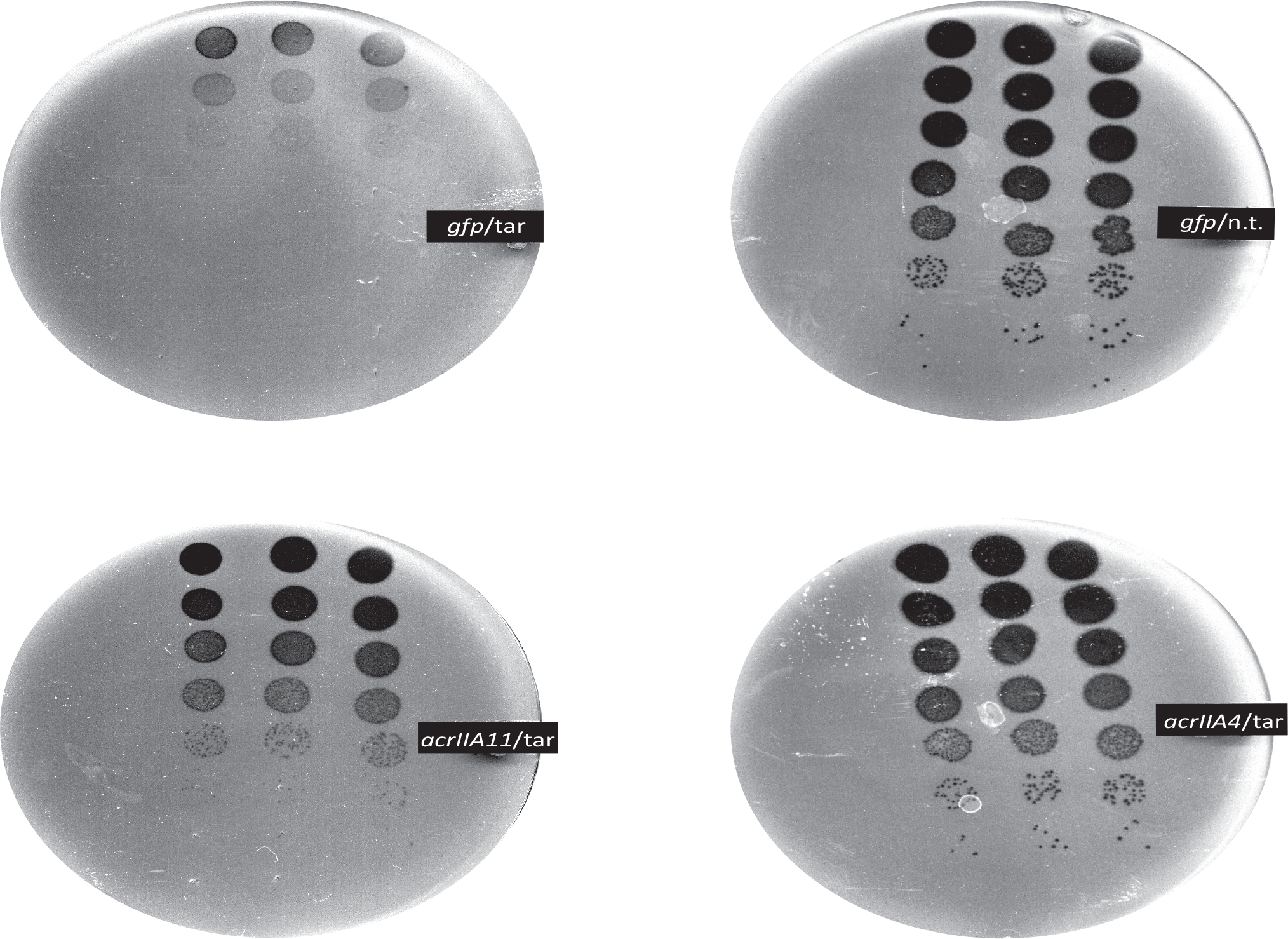
*AcrIIA11* protects phage from SpyCas9. Mu phage fitness, measured by plaquing on *E. coli* expressing Mu-targeting SpyCas9, is measured in the presence of *gfp*, *acrIIA11*, or *acrIIA4* via serial ten-fold dilutions. Bacterial clearing (black) occurs when phage Mu overcomes Cas9 immunity and lyses *E. coli*. Based on a non-targeting (n.t.) crRNA control, we conclude that SpyCas9 confers 10^5^-fold protection against phage Mu in these conditions. Both *acrIIA11* and *acrIIA4* significantly enhance Mu fitness by inhibiting SpyCas9. The indicated anti-CRISPR gene or *gfp* control is expressed from a second plasmid, in trans.

**Figure S7.**
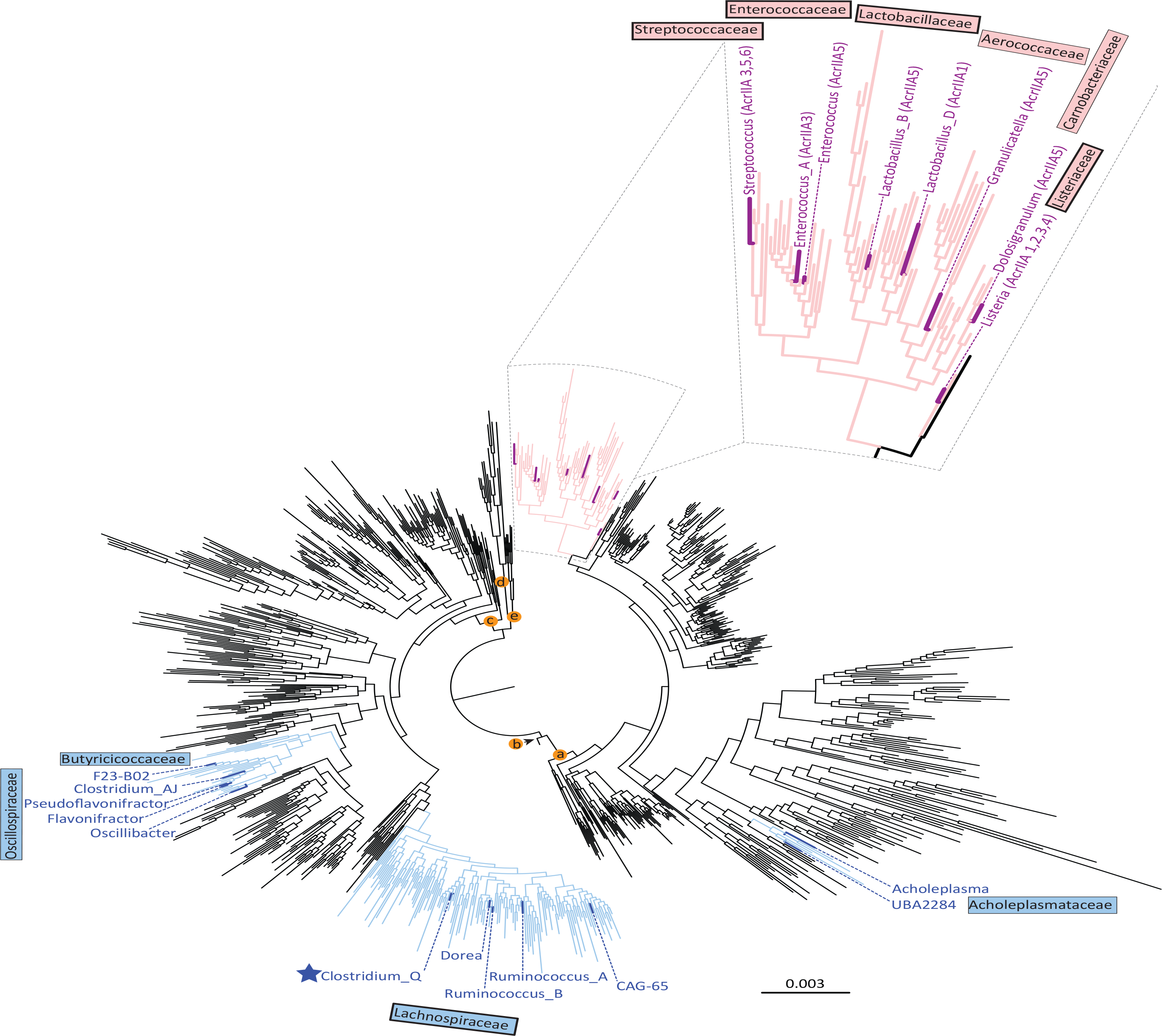
*AcrIIA11* is widely distributed across Firmicutes and related phyla compared to *acrIIA1-acrIIA6*. As in figure 3D, each node represents a genus and families colored light blue contain *acrIIA11* homologs while families colored pink contain homologs of *acrIIA1* - *acrIIA6*. Genera are colored dark blue or magenta if *acr* homologs could confidently be assigned to this phylogenetic resolution. The star indicates the *Clostridium_Q* genus depicted in figure 2C. A thick border around a family indicates that is enriched for type II-A CRISPR-Cas systems relative to all bacterial families (chi-squared test, p<1×10^−4^, family classification used the GTDB scheme with functional attributions provided by AnnoTree). We used the GTDB taxonomy because it helps to resolve a well-recognized polyphyly among *Clostridia* and ensures that taxonomic labels are linked to monophyletic groups while applying taxonomic ranks (phylum, class, order, etc.) at even phylogenetic depths. Lettered nodes indicate GTDB phyla (a: Firmicutes, b: Firmicutes_D, c: Firmicutes_A, d: Fusobacteria, e: Firmicutes_F).

**Figure S8.**
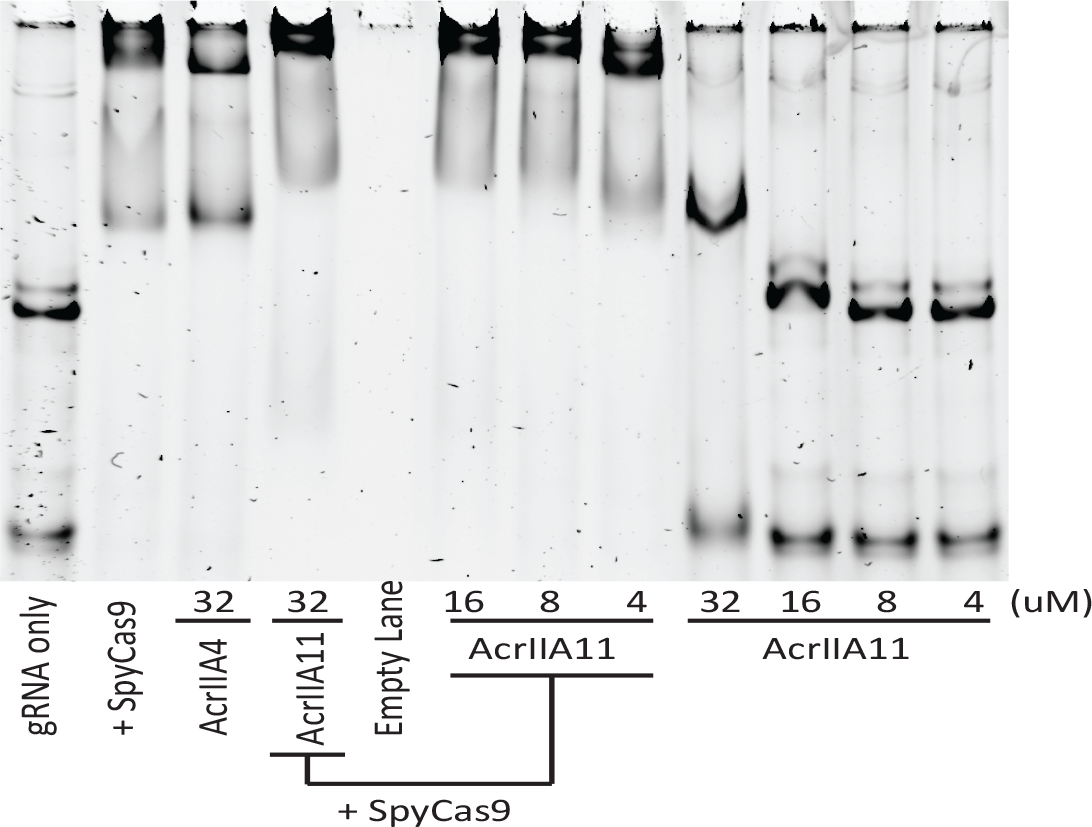
AcrIIA11 does not prevent a gRNA gel-shift, indicating that the RNA is protein-bound. An EMSA examining the relative mobility of *S. pyogenes* gRNA (0.2µM) through an 8% acrylamide gel in the presence of SpyCas9 and/or various Acrs. Neither AcrIIA4 nor AcrIIA11 prevent a gel-shift shift upon SpyCas9 addition, though the nature of the shift is different between Acrs. AcrIIA11 also appears to super-shift the SpyCas9/gRNA complex, which may represent AcrIIA11 bound to this complex. At high concentrations, AcrIIA11 binds gRNA.

**Figure S9.**
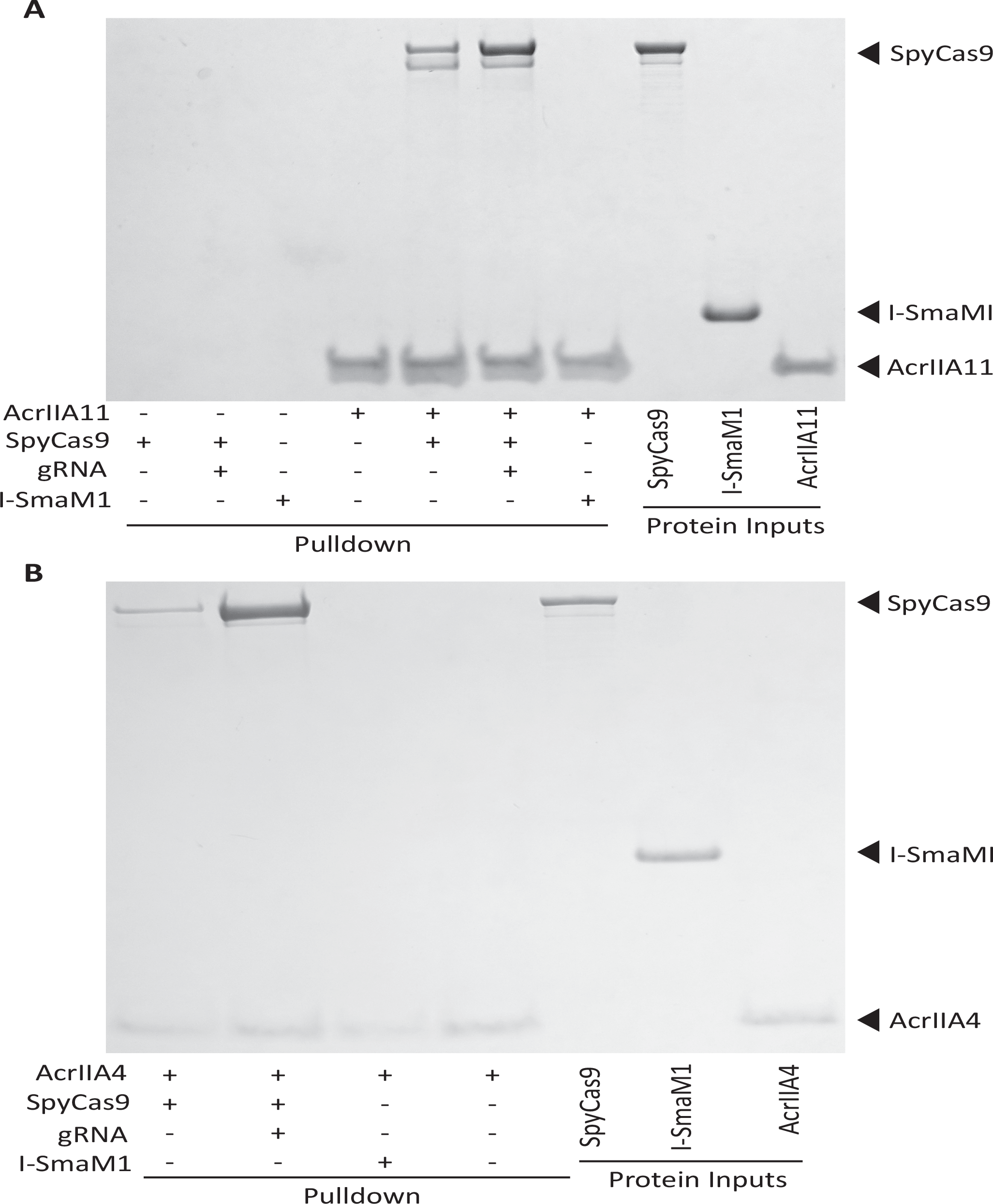
AcrIIA11 binds SpyCas9 with a moderate preference for the gRNA-loaded form. (**A)** AcrIIA11 binds SpyCas9. **(B)** AcrIIA4 binds SpyCas9. SpyCas9 and gRNA were pre-incubated before mixing with a 2x-strep-tagged AcrIIA11 (A) or AcrIIA4 (B). SpyCas9 without gRNA and the meganuclease I-SmaMI were also used. (A) Pulldowns on AcrIIA11 brought with them SpyCas9 but not I-SmaMI, and the presence of gRNA improved the strength of this interaction, but not to the degree seen with AcrIIA4 in (B). These images depict total protein content visualized by Coomassie stain. The gel in (A) is identical to that depicted in fig. 5D except that the three leftmost control lanes have not been cropped from this image.

**Table S1.** False positive genotypes and frequencies encountered during the development of the Cas9 Acr selection. Asterisks (*) indicate genotyped experiments that are referenced in figure S2. Parentheses indicate the number of colonies queried and the number with the indicated genotype.

| Experiment | Target Plasmid | pCas9 used (table S5) | Cas9 expression | No. of target sites | Iteration No. | pCas9 genotype | Target site genotype | Surviving Proportion of Kan<sup>R</sup> Transformants | Notes |
| --- | --- | --- | --- | --- | --- | --- | --- | --- | --- |
| A | pZE21-GFP | pCas9_crA | Constitutive | 1 | 1 | loss-of-function mutant (8/8) | wild-type (8/8) | not determined | Constitutive Cas9 restricted transformation |
| B | pZE21-GFP | pCas9_crZ | Constitutive | 1 | 1 | loss-of-function mutant (6/6) | wild-type (6/6) | not determined | Constitutive Cas9 restricted transformation |
| C* | pZE21-GFP | pCas9_in_crA | Inducible | 1 | 1 | loss-of-function mutant (6/6) | wild-type (6/6) | 5.05E-06 | In figure S2 |
| D | pZE21-GFP | pCas9_in_crA | Inducible | 1 | 1 | no colonies tested | no colonies tested | 1.87E-05 | In figure S2 |
| E* | pZE21-GFP | pCas9_in_crA | Inducible | 1 | 2 | wild-type (6/6) | PAM or protospacer mutant (12/12) | 0.03 | In figure S2 |
| F | pZE21-GFP | pCas9_in_crA | Inducible | 1 | 1 | no colonies tested | no colonies tested | 4.82E-05 | In figure S2 |
| G* | pZE21-GFP | pCas9_in_crAcrZ | Inducible | 2 | 1 | loss-of-function mutant (11/11) | wild-type (10/10) | 7.03E-06 | In figure S2 |
| H | pZE21-GFP | pCas9_in_crA | Inducible | 1 | 1 | no colonies tested | no colonies tested | 2.98E-05 | In figure S2 |
| I | pZE21-GFP | pCas9_in_crAcrZ | Inducible | 2 | 1 | no colonies tested | no colonies tested | 1.06E-05 | In figure S2 |
| J* | pZE21-GFP | pCas9_in_crA | Inducible | 1 | 2 | loss-of-function mutant (4/4) | wild-type (8/8) | 2.72E-05 | In figure S2 |
| K* | pZE21-GFP | pCas9_in_crAcrZ | Inducible | 2 | 2 | loss-of-function mutant (6/6) | wild-type (9/9) | 3.44E-06 | In figure S2 |
| L* | pZE21-GFP | pCas9_in_crAcrZ | Inducible | 2 | 2 | loss-of-function mutant (6/6) | wild-type (20/20) | 1.46E-05 | In figure S2 |
| M | pZE21-GFP | pCas9_in_crAcrZ | Inducible | 2 | 1 | no colonies tested | no colonies tested | 4.20E-05 | In figure S2 |
| N | pZE21-GFP | pCas9_in_crAcrZ | Inducible | 2 | 2 | no colonies tested | no colonies tested | 3.77E-04 | In figure S2 |
| O | pZE21-GFP | pCas9_in_crAcrB | Inducible | 2 | 1 | no colonies tested | no colonies tested | 3.75E-05 | In figure S2 |
| P | pZE21-GFP | pCas9_in_crAcrB | Inducible | 2 | 2 | no colonies tested | no colonies tested | 1.27E-04 | In figure S2 |

**Table S2.** Metagenomic library information. Each library has an average DNA insert size of ~2 Kb (Clemente et al., 2015; Pehrsson et al., 2016). In calculating the number of clones subjected to Cas9 selection, we assumed that transformation is a Poisson process.

| Library Information |  | Iteration #1 |  |  | Iteration #2 |  |  |
| --- | --- | --- | --- | --- | --- | --- | --- |
| Library | Unique Clones (x10<sup>6</sup>) | Transformants (x10<sup>6</sup>) | Unique Clones Subjected to Cas9 Selection (x10<sup>6</sup>) | KanR Colonies Collected | Amt. Transformed (ng) | Transformants (x10<sup>6</sup>) | Titer Volumes Collected |
| Oral_3 | 3.65 | 5 | 2.72 | 2850 | 272 | 56 | 2ml / 100µl |
| Oral_5 | 3.88 | 7.85 | 3.37 | 3580 | 396 | 71.8 | 100µl / 1µl |
| Fecal_01A | 1.65 | 21.4 | 1.65 | 3110 | 261 | 32.9 | 1µl / 0.01µl |
| Fecal_01E | 1.35 | 4.88 | 1.31 | 520 | 236 | 29.8 | 100µl / 1µl |
| Fecal_03G | 2.7 | 4.36 | 2.16 | 2910 | 216 | 46.2 | 1µl / 0.01µl |
| GFP | n/a | 42.8 | n/a | 5650 | 308 | 3.86 | 100µl |

**Table S3.**
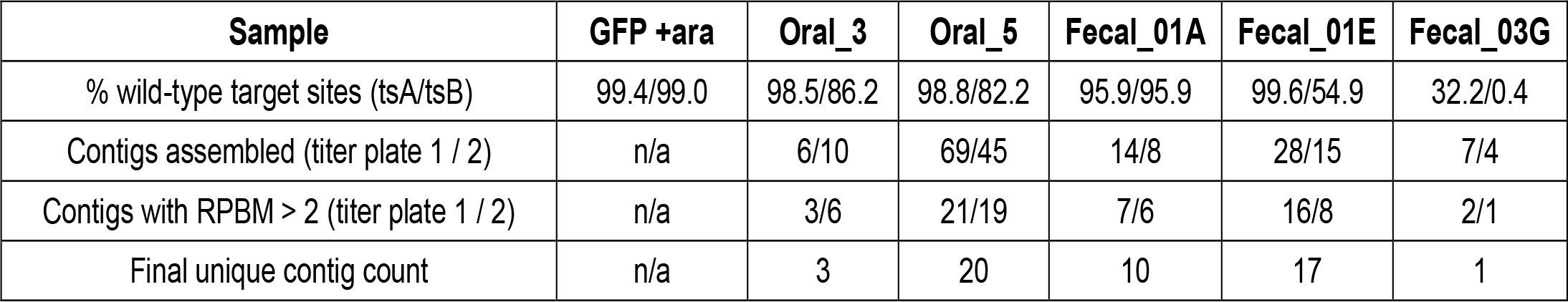
Summary of twice-iterated functional selections for Cas9 Acrs. Wild-type target site statistics were determined using bulk plasmids surviving two rounds of SpyCas9 selection. The ‘ts’ acronym stands for ‘target site’, which encompasses both the protospacer and PAM sequences required for SpyCas9 restriction.

| **Sample** | **GFP +ara** | **Oral_3** | **Oral_5** | **Fecal_01A** | **Fecal_01E** | **Fecal_03G** |
| --- | --- | --- | --- | --- | --- | --- |
| % wild-type target sites (tsA/tsB) | 99.4/99.0 | 98.5/86.2 | 98.8/82.2 | 95.9/95.9 | 99.6/54.9 | 32.2/0.4 |
| Contigs assembled (titer plate 1 / 2) | n/a | 6/10 | 69/45 | 14/8 | 28/15 | 7/4 |
| Contigs with RPBMs > 2 (titer plate 1 / 2) | n/a | 3/6 | 21/19 | 7/6 | 16/8 | 2/1 |
| Final unique contig count | n/a | 3 | 20 | 10 | 17 | 1 |

**Table S4.**
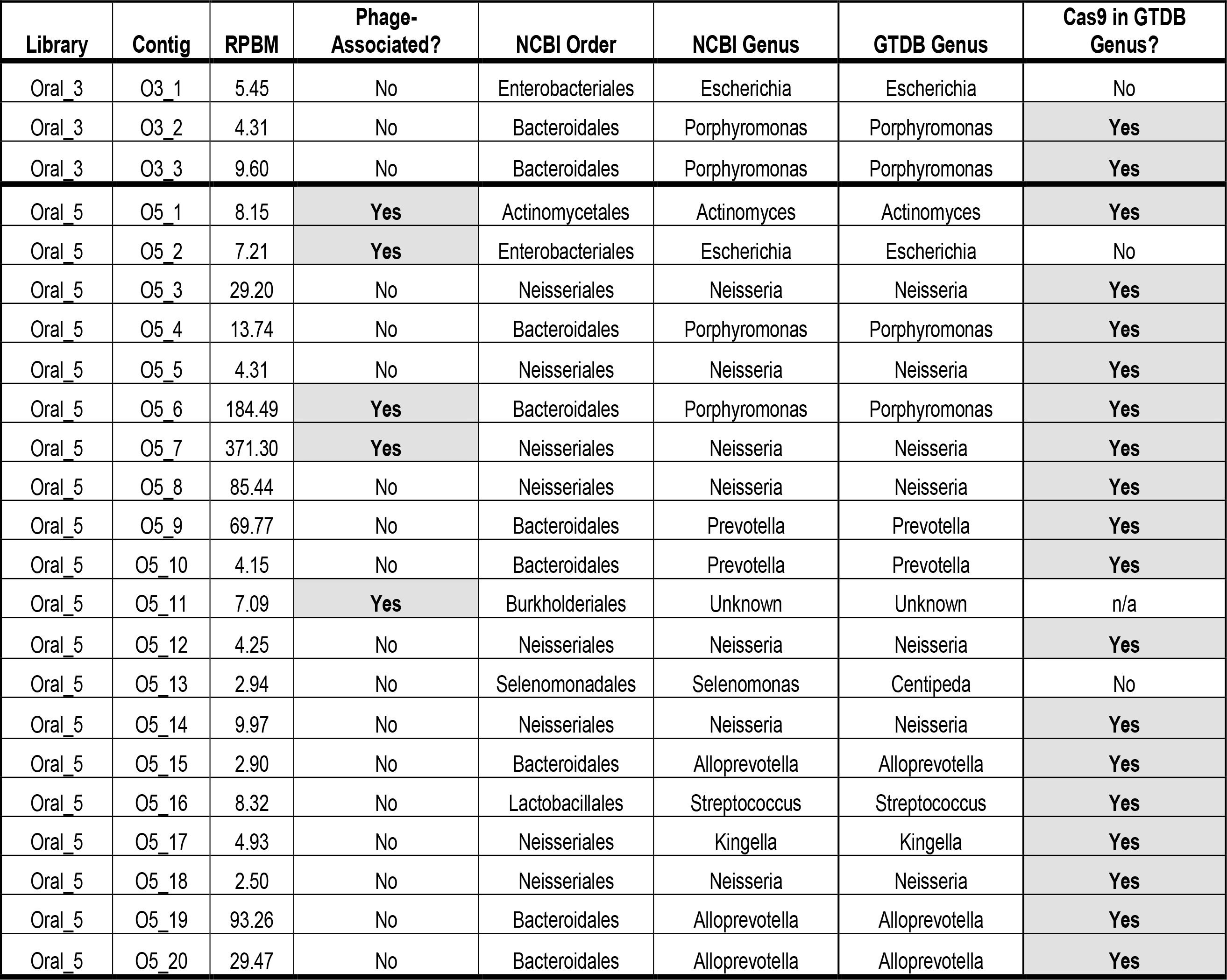
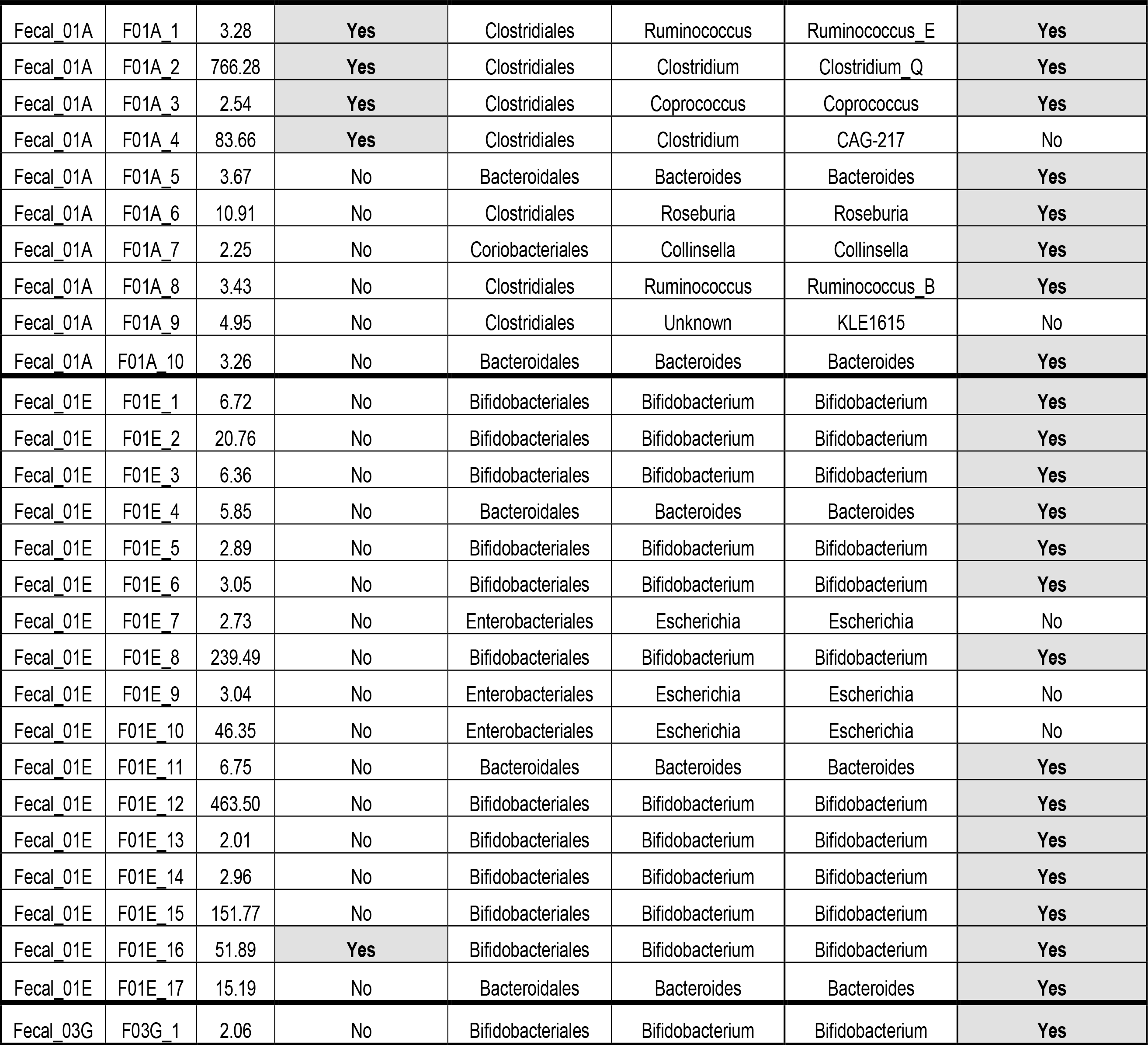
All contigs in the final dataset. Contigs are organized by library. Reads per base-pair per million reads (RPBM) indicate coverage for a given contig, which is a proxy for that sequence’s abundance following SpyCas9 selection. Proteins with close homologs in a predicted phage were used to classify a given contig as phage-associated (see methods). NCBI taxa were determined by blastn and blastp searches of contig and protein sequences; GTDB taxa were determined as described in the methods and were used throughout most of the manuscript to resolve a well-recognized polyphyly among the Clostridiales.

**Table S5.**
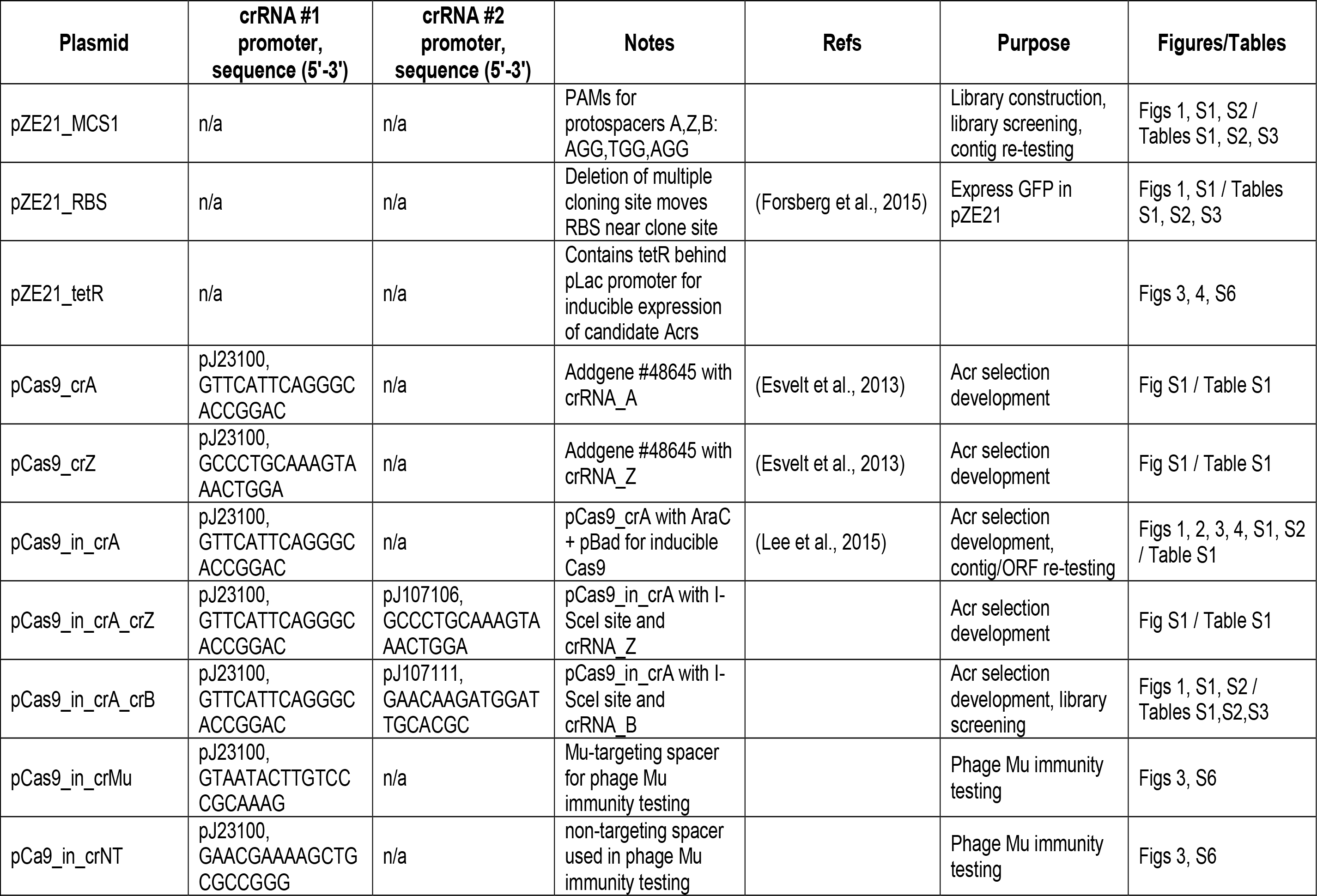
Plasmids used in this study.

**Table S6.**
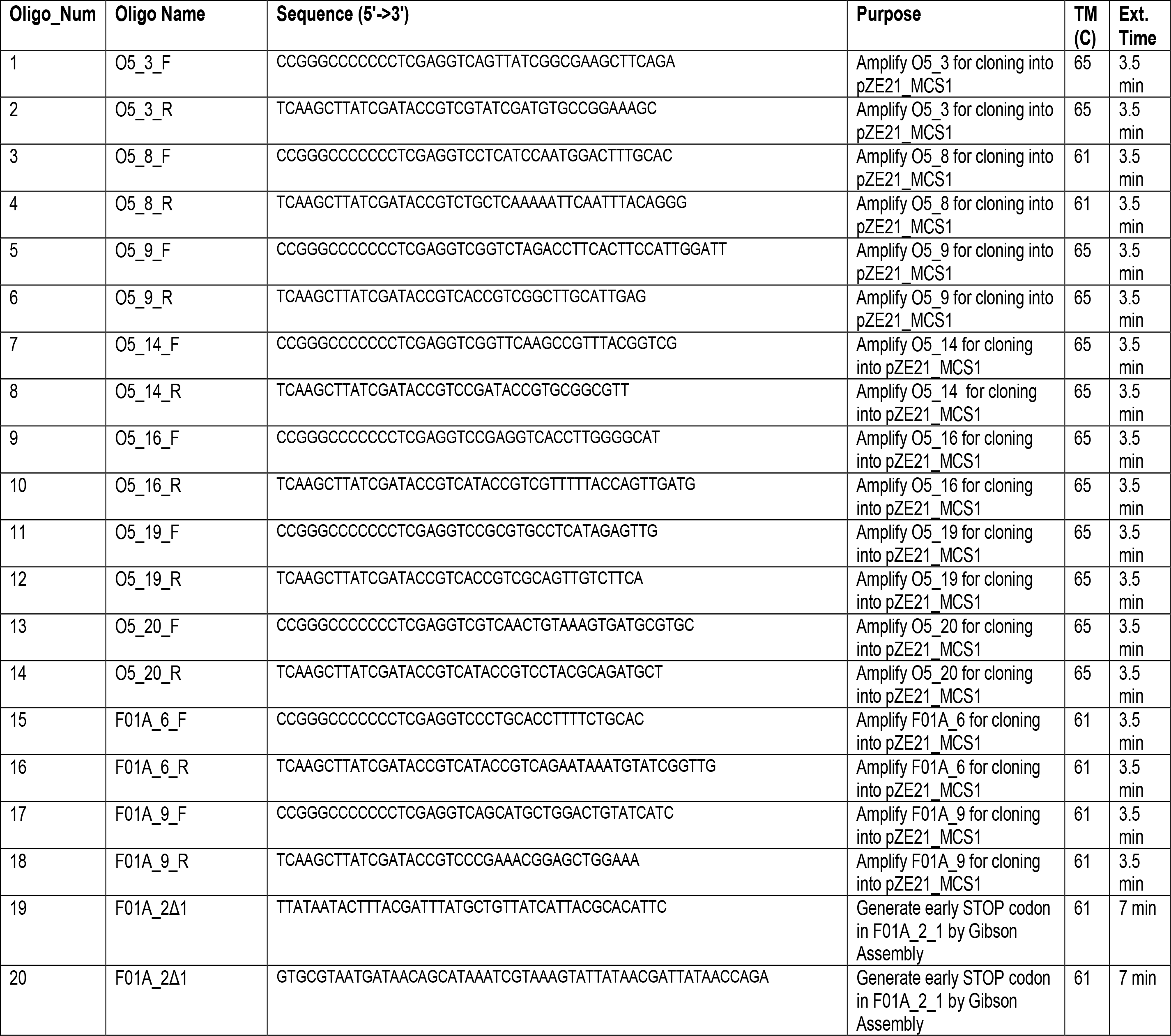
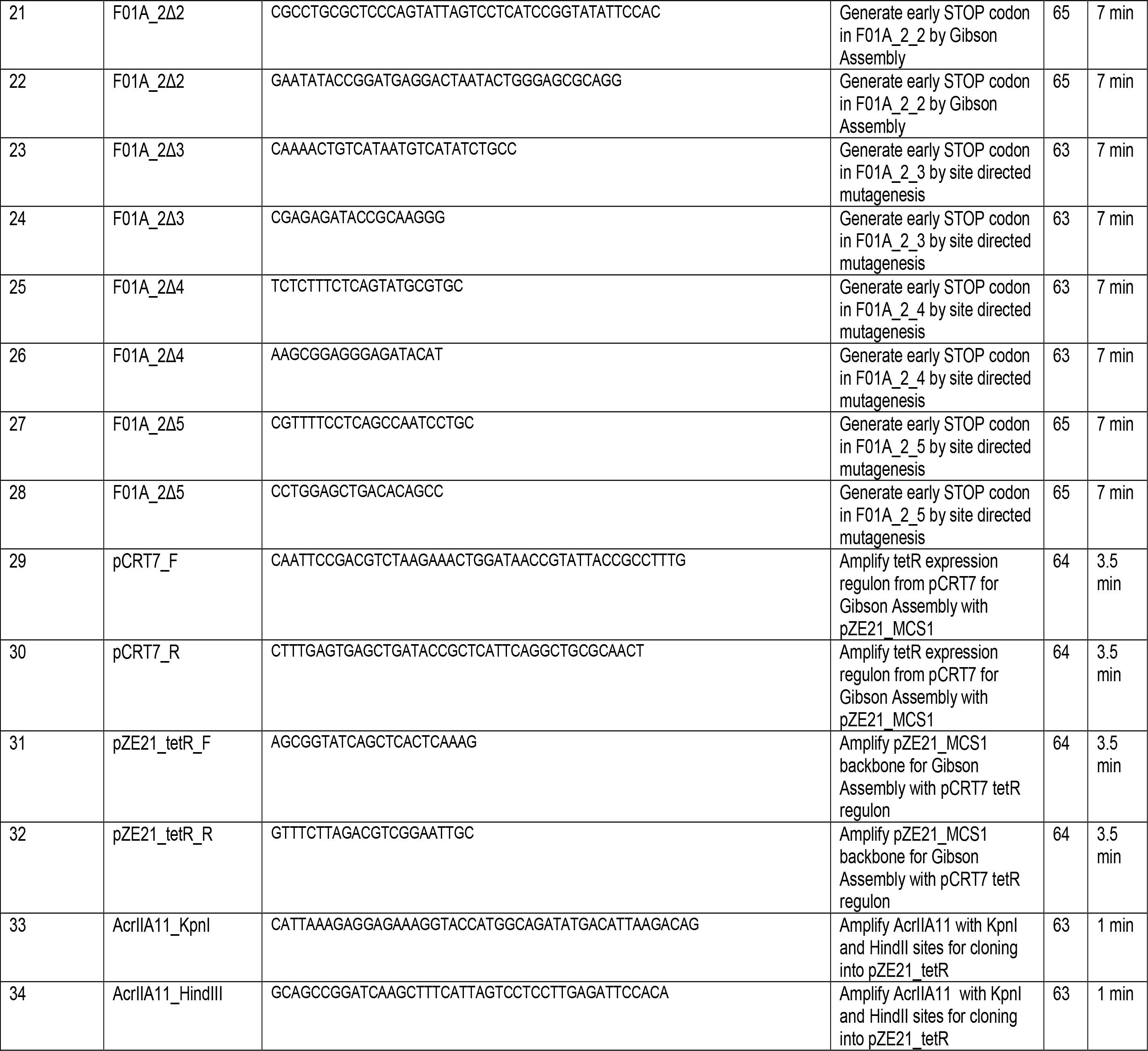
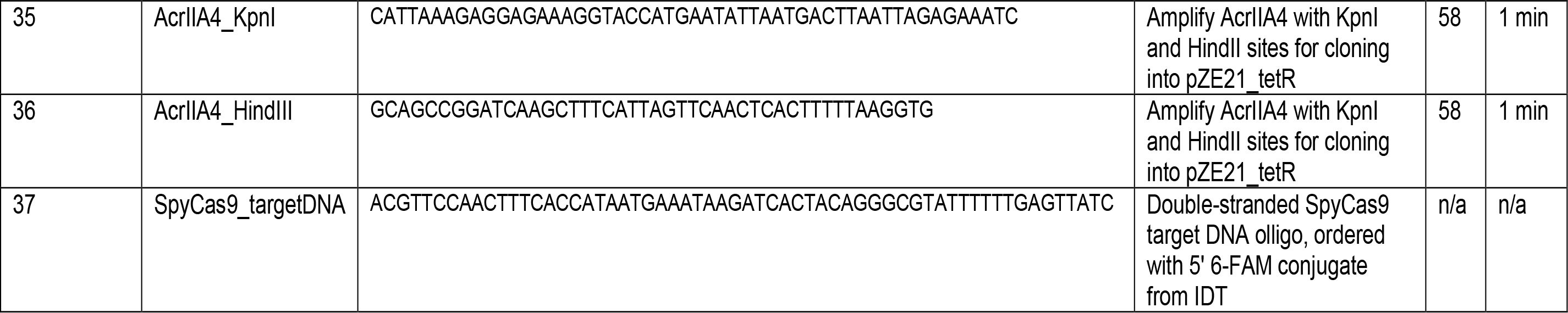
Oligos used in this study.

**Table S7.**
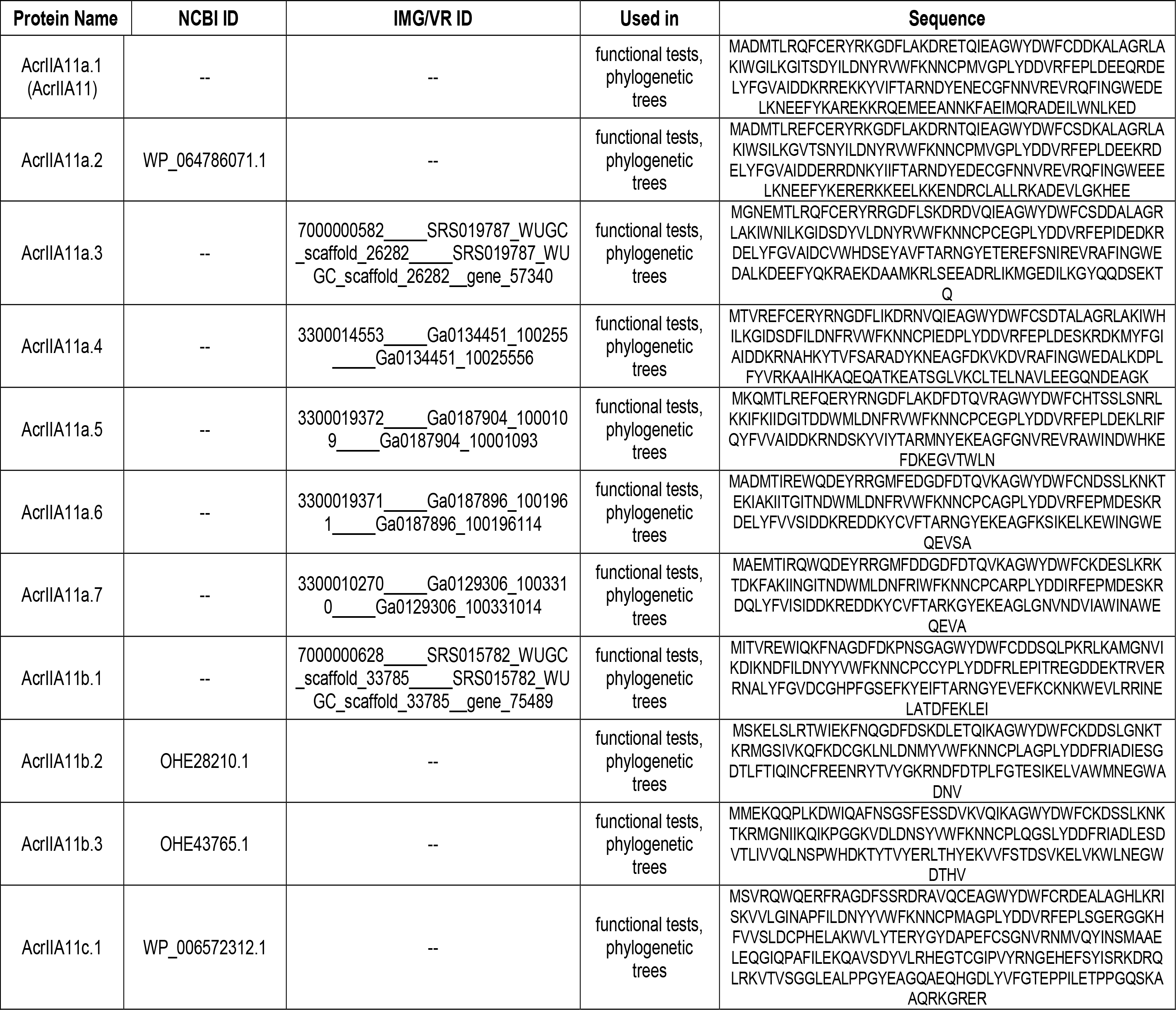
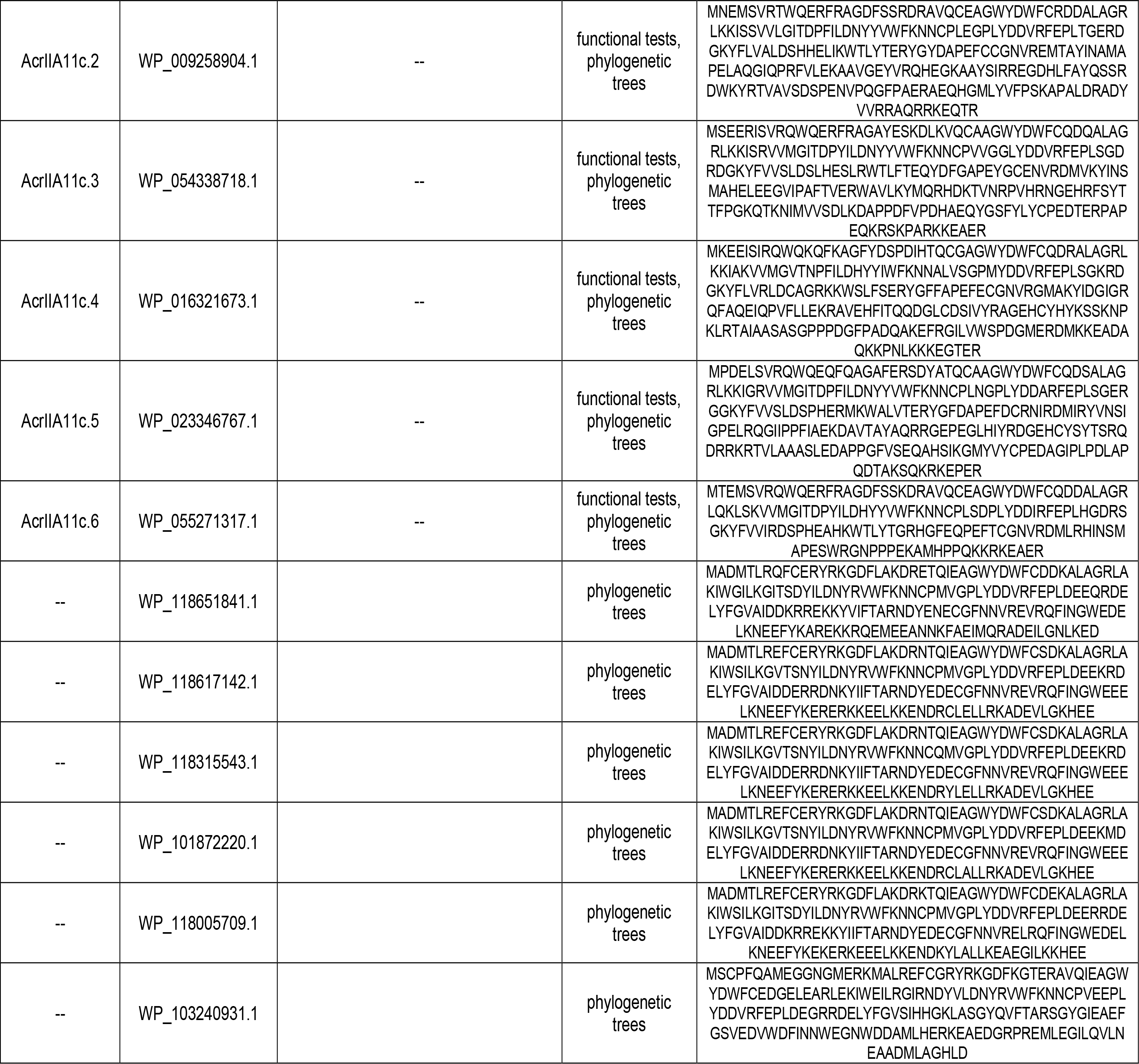
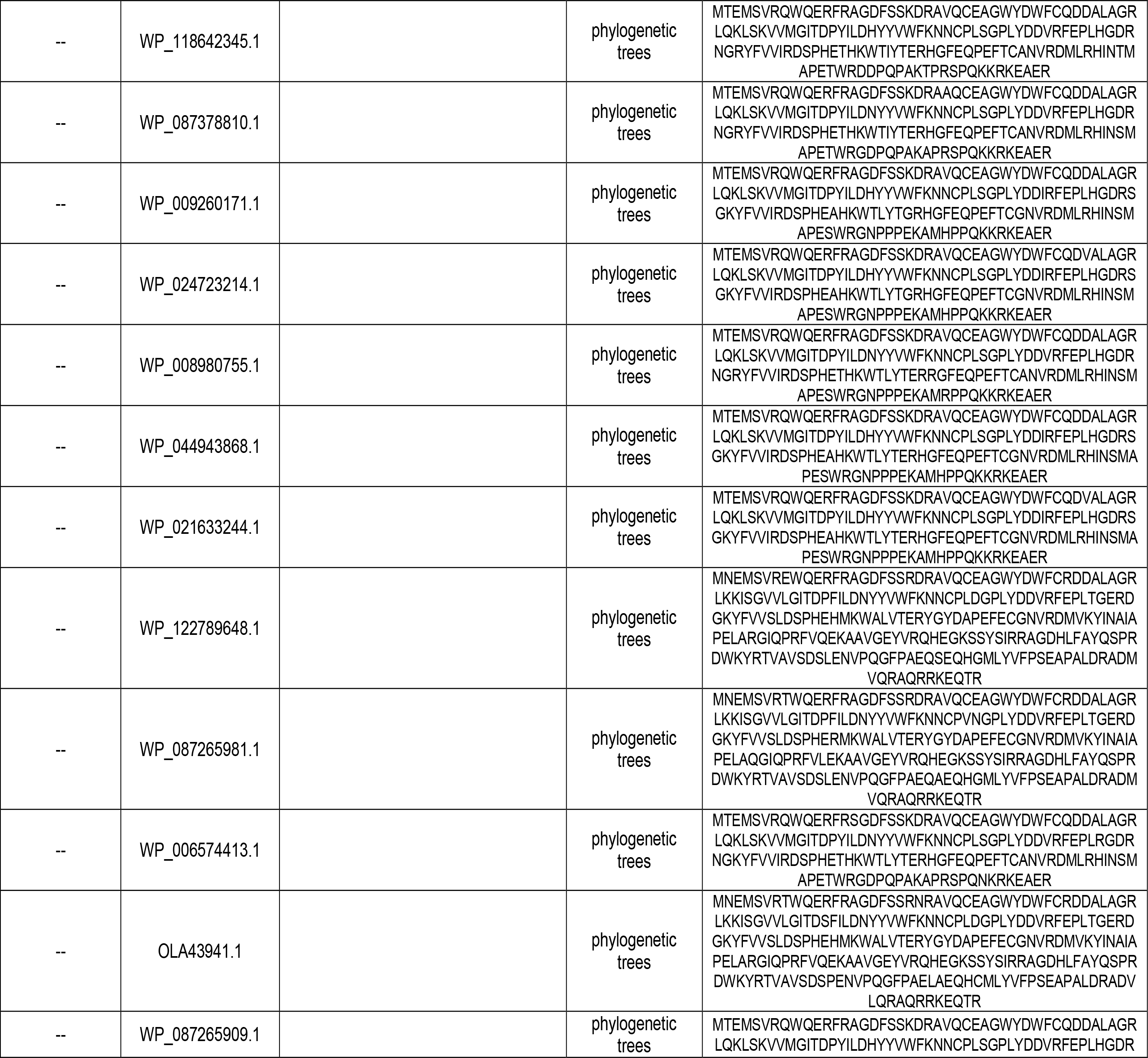
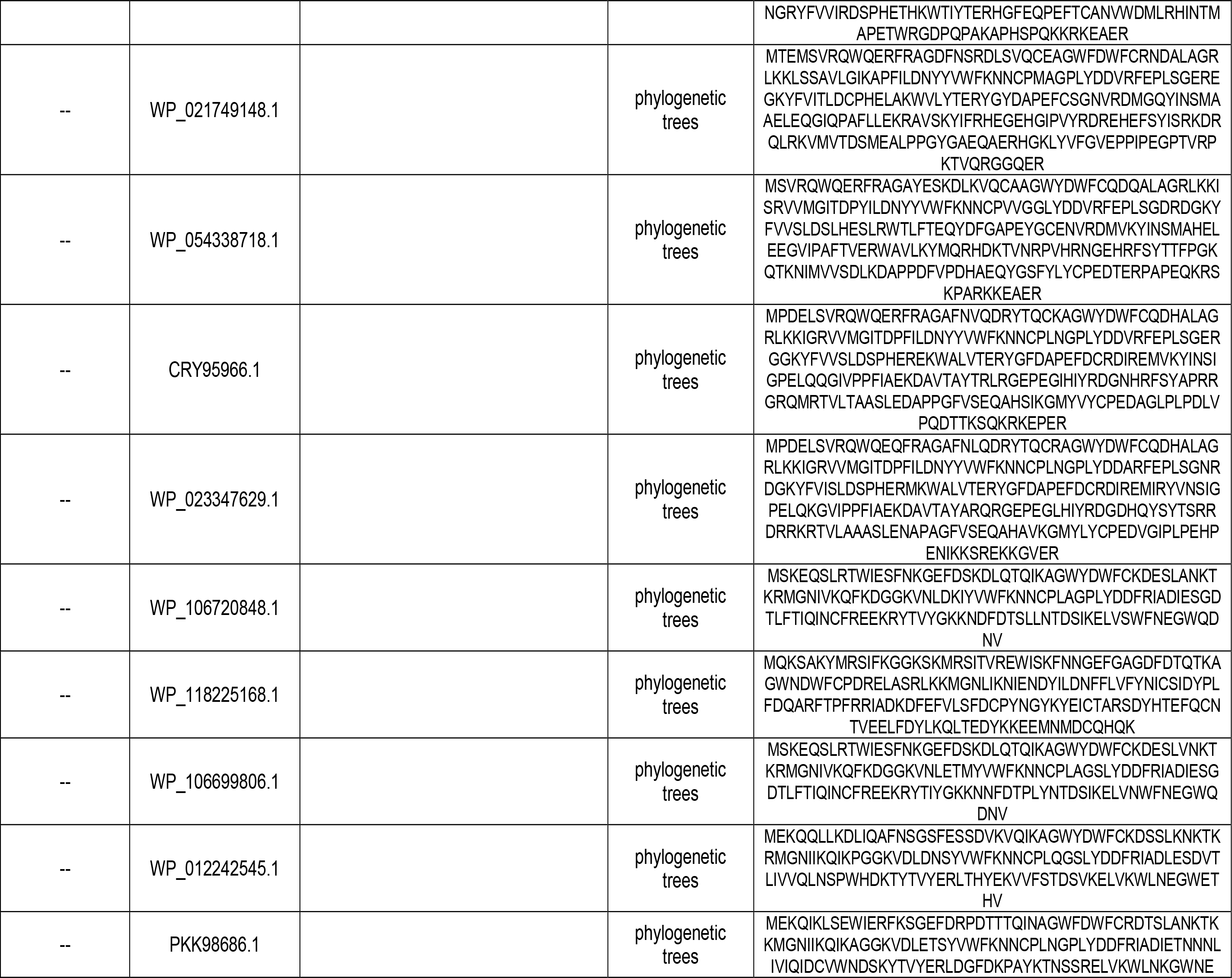
Acr sequences used to create the phylogenetic tree in figure 4A.

